# A conserved mechanism for meiotic chromosome organization through self-assembly of a filamentous chromosome axis core

**DOI:** 10.1101/375220

**Authors:** Alan M.V. West, Scott C. Rosenberg, Sarah N. Ur, Madison K. Lehmer, Qiaozhen Ye, Götz Hagemann, Iracema Caballero, Isabel Usón, Franz Herzog, Kevin D. Corbett

## Abstract

The meiotic chromosome axis plays key roles in meiotic chromosome organization and recombination, yet the underlying protein components of this structure are highly diverged. Here, we show that “axis core proteins” from budding yeast (Red1), mammals (SYCP2/SYCP3), and plants (ASY3/ASY4) are evolutionarily related and play equivalent roles in chromosome axis assembly. We first identify motifs in each complex that recruit meiotic HORMADs, the master regulators of meiotic recombination. We next find that axis core complexes form homotetrameric (Red1) or heterotetrameric (SYCP2:SYCP3 and ASY3:ASY4) coiled-coil assemblies that further oligomerize into micron-length filaments. Thus, the meiotic chromosome axis core in fungi, mammals, and plants shares a common molecular architecture and role in axis assembly and recombination control. We propose that the meiotic chromosome axis self-assembles through cooperative interactions between dynamic DNA loop-extruding cohesin complexes and the filamentous axis core, then serves as a platform for chromosome organization, recombination, and synaptonemal complex assembly.

## Introduction

Meiosis is a specialized cell division program that generates haploid gametes from a diploid cell, in preparation for sexual reproduction. Meiosis achieves a two-fold reduction in ploidy through two successive cell divisions without an intervening DNA replication step. Homologous chromosomes segregate from one another in the first meiotic division (meiosis I), and replicated sister chromosomes segregate in meiosis II. Accurate segregation of homologs in meiosis I requires that homologs identify and physically link to one another in the extended meiotic prophase. Homolog recognition and physical association is achieved through crossover formation, in which programmed double strand DNA breaks (DSBs) in each chromosome are repaired in a specialized homologous recombination pathway, resulting in a reciprocal exchange of genetic information and the physical linkage of homologs.

A highly-conserved meiosis-specific structure, the chromosome axis, assembles in early meiotic prophase and provides a scaffold for the organization of chromosome as a linear array of loops (1-3), and orchestrates the formation of DSBs and their repair as inter-homolog crossovers (4-14). Components of the chromosome axis include DNA-binding and -organizing cohesin complexes (15), plus proteins of the meiotic HORMAD family that regulate DSB and crossover formation (16-24). Most organisms also possess additional factors, here termed “axis core” proteins, that are important for axis formation and meiotic HORMAD recruitment. The archetypal axis care protein is *S. cerevisiae* Red1, an 827-residue protein that recruits the HORMAD protein Hop1 to the axis via a HORMA domain-binding “closure motif’ (25, 26). A conserved region at the Red1 C-terminus is predicted to adopt a coiled-coil structure and mediates self-association of the protein (26, 27), suggesting that oligomer formation by Red1 may be important for axis function.

While clearly-identifiable Red1 homologs do not exist outside fungi, many other organisms possess functionally-equivalent axis core proteins with predicted coiled-coil structure. Mammals possess two such proteins, SYCP2 (1500 residues in *Mus musculus*) and SYCP3 (254 residues), which are both required for proper axis formation and wild-type levels of crossovers (28, 29), and are known to interact with one another through their C-terminal coiled-coil domains (30). SYCP2 and SYCP3 are interdependent for their axis localization (28, 30-32), and a mutant of SYCP2 lacking its C-terminal coiled-coil region shows a loss of SYCP3 from the axis (30, 32), suggesting that the SYCP2 N-terminal region mediates localization of the complex, while the C-terminal domain mediates oligomerization with SYCP3. In plants, the axis proteins ASY3 (793 residues in *Arabidopsis thaliana*) and ASY4 (212 residues) proteins are both important for crossover formation, and these proteins also associate with one another through their C-terminal coiled coil domains (33, 34). While neither SYCP2/SYCP3 nor ASY3/ASY4 has been reported to possess HORMAD-interacting closure motifs, the similar domain structure and roles in crossover formation between these protein families and *S. cerevisiae* Red1 suggests that they may be evolutionarily related (33, 35).

In addition to its roles in chromosome organization and crossover formation in early meiotic prophase, the chromosome axis plays a later role as a key structural element of the highly-conserved yet functionally enigmatic synaptonemal complex (SC). As inter-homolog crossovers form, the chromosome axes of each homolog pair, now termed “lateral elements” of the SC, become linked by coiled-coil “transverse filaments” along their entire length (19, 36-40). In fungi, plants, and mammals, SC assembly is tightly coordinated with removal of the meiotic HORMADs from the chromosome axis by the AAA+-ATPase Pch2/TRIP13, in a key feedback mechanism controlling crossover levels (37, 38, 40-42). SC assembly is required for crossover maturation, and serves as a signal to the cell that a given homolog pair has obtained crossovers (1, 43).

While the molecular architecture of the SC transverse filaments and associated “central element” is becoming better understood (44-48), the architecture of the chromosome axis/SC lateral element has remained largely uncharacterized. Specifically, it is not known whether mammalian and plant axis core proteins possess HORMAD-binding closure motifs like Red1, leaving open the question of how HORMADs are recruited to chromosomes in these organisms. More significantly, the oligomeric structure of the axis core proteins, whether this structure is conserved, and how this structure contributes to the axis’s roles in chromosome organization, inter-homolog recombination, and SC architecture are important open questions. Mammalian SYCP3 is known to form coiled-coil homotetramers (49) that self-associate into larger structures that can be observed both in cell culture (31, 50) and in vitro (49, 51, 52), but how SYCP3 cooperates with SYCP2 to mediate chromosome localization and axis assembly is not known. Neither fungal Red1 nor plant ASY3/ASY4 has been characterized biochemically, leaving open the question of how these proteins self-assemble, and whether these assemblies resemble those of mammalian SYCP3.

Here, we address these questions and establish that the molecular architecture of the meiotic chromosome axis is shared between fungi, mammals and plants. We find that budding-yeast Red1 forms stable homotetrameric complexes via its coiled-coil C-terminus, and that these tetramers associate end-to-end to form extended filaments visible by electron microscopy. We identify HORMAD-binding closure motifs in both mammalian SYCP2 and plant ASY3, supporting these proteins’ identification as Red1 homologs and strongly suggesting a role in meiotic HORMAD recruitment to meiotic chromosomes. We further show that both SYCP2/SYCP3 and ASY3/ASY4 form heterotetrameric coiled-coil complexes that self-assemble into extended filaments, paralleling our findings with Red1. Taken together, these data reveal common principles of meiotic chromosome axis assembly and function that are widely shared throughout eukaryotes.

## Results

### Budding Red1 forms filaments from coiled-coil tetramer units

In budding yeast, the chromosome axis is made up of the HORMAD protein Hop1, its binding partner Red1, and cohesin complexes containing the meiosis-specific Rec8 kleisin subunit (1, 53). We and others have outlined the assembly mechanisms of Hop1, which binds short “closure motifs” in its own C-terminal tail and in Red1 through its conserved HORMA domain (Figure 1A) (25, 26). Red1 is less well-understood. This protein possesses a conserved N-terminal domain immediately followed by a Hop1-binding closure motif, an extended linker domain with high predicted disorder, and a C-terminal domain that mediates Red1 self-association and is predicted to adopt a coiled-coil structure (Figure 1A) (25-27). Prior genetic studies isolated two point-mutations in the Red1 C-terminal domain, I743A (54) and I758R (55), that each strongly affect both SC assembly and spore viability in *S. cerevisiae*. While these phenotypes were attributed to effects on binding other meiotic chromosome-associated proteins, these residues’ location within a predicted coiled-coil domain prompted us to consider instead that the observed defects in these mutants may due to a disruption of a Red1 oligomer important for meiotic chromosome axis function.

**Figure 1.**
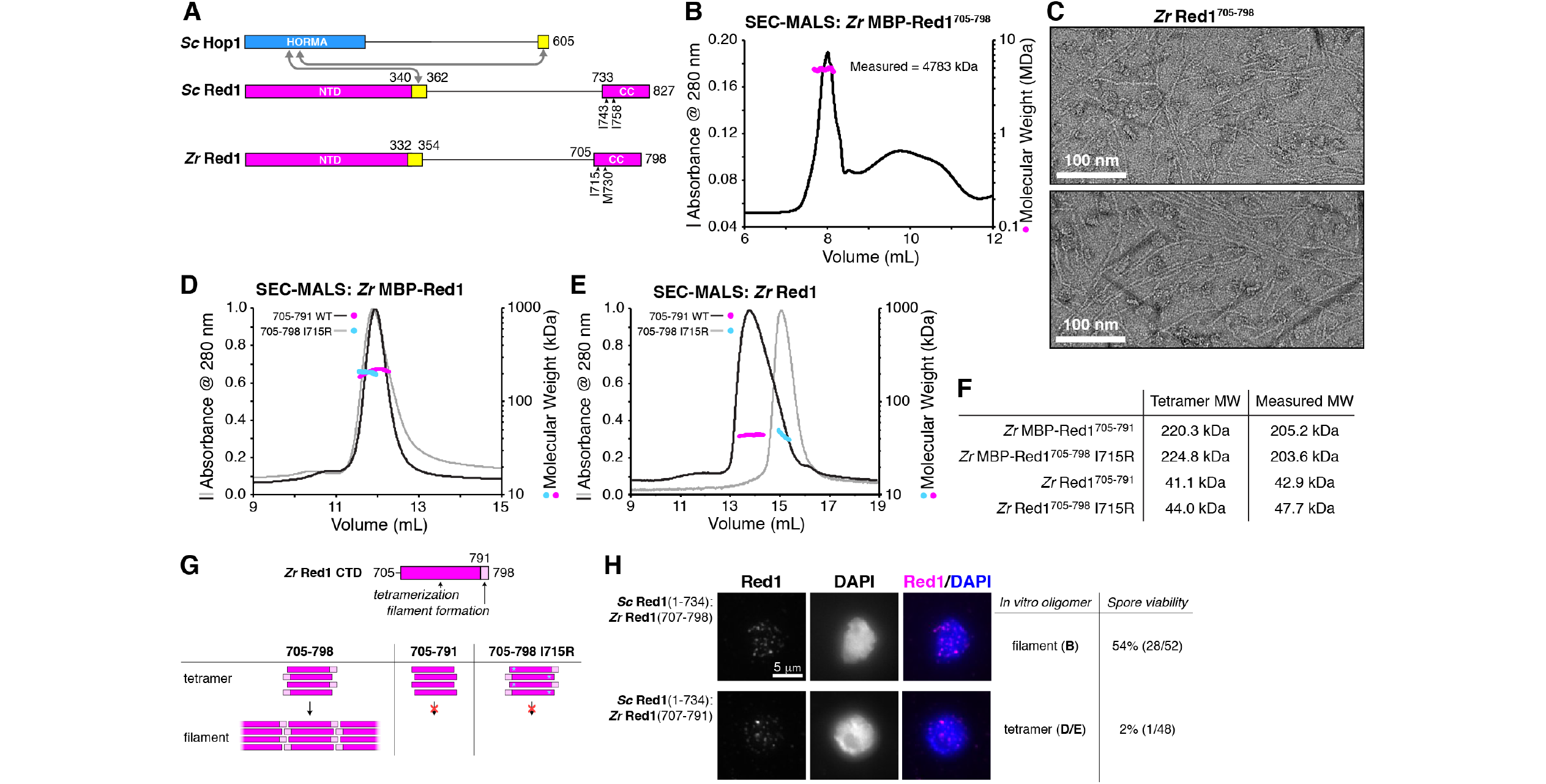
Molecular architecture of the budding-yeast Red1 C-terminal domain. (A) Schematic of *S. cerevisiae* (*Sc*) chromosome axis proteins Hop1 and Red1, and *Zygosaccharocmyces zouxii* (*Zr*) Red1. Yellow regions indicate Hop1-binding closure motifs (25). For *S. cerevisiae* Red1, the positions of two previously-identified mutations in the C-terminal domain that disrupt axis function, I743R (54) and I758R (55), are shown. See Figure 1 – Figure Supplement 1 for sequence alignment of the Red1 C-terminal domain. (B) SEC-MALS analysis of purified His_6_-MBP-*Zr* Red1^705-798^. Calculated molecular weight of a monomer = 55.9 kDa; Measured molecular weight = 4783 kDa (~85-mer). (C) Representative negative-stain electron micrographs of purified untagged *Zr* Red1^705-798^. See Figure 1 – Figure Supplement 2A for additional full micrographs, and Figure 1 – Figure Supplement 2B for micrographs of His_6_-MBP-*Zr* Red1^705-798^. (D) SEC-MALS analysis of purified His_6_-MBP-*Zr* Red1^705-791^ and His_6_-MBP-*Zr* Red1^705-791^ I715R. (E) SEC-MALS analysis of purified *Zr* Red1^705-791^ and *Zr* Red1^705-791^ (F) Table summarizing SEC-MALS results from (D) and (E). (G) Schematic of *Zr* Red1 C-terminal domain oligomerization. Wild-type *Zr* Red1^705-798^ forms homotetramers that further oligomerize into extended filaments. Removal of the C-terminal seven amino acids (Zr Red1^705-791^) or mutations of M715 to arginine (Zr Red1^705-598^ M715R) results in loss of filament formation but maintenance of tetramer formation. (H) Spore viability and meiotic prophase chromosome localization of *S. cerevisiae/Z. rouxii* Red1 chimeric constructs, imaged at three hours after meiotic induction. Removal of the C-terminal seven residues of *Z. rouxii* Red1 from the chimera (which eliminates filament formation in vitro) does not affect chromosome localization, but dramatically affects spore viability.

To test this idea, we expressed in *E. coli*, and purified the Red1 C-terminal domain from several budding yeasts, and found that uniformly, these proteins formed large assemblies as measured by size-exclusion chromatography (Figure 1B and data not shown). We examined one Red1 construct, *Zygosaccharomyces rouxii (Zr*) Red1^705-798^, by negative-stain electron microscopy. We observed filaments up to several microns in length (Figure 1C), suggesting that the large assemblies of purified Red1 C-terminal domain are not disordered aggregates but rather represent a biologically relevant structure. In the course of construct optimization, we also cloned and purified a truncated *Zr* Red1 construct missing the C-terminal seven residues of the protein (Zr Red1^705-791^). Strikingly, this construct did not form assemblies in solution, but rather formed stable homotetramers as measured by size-exclusion chromatography coupled to multiangle light scattering (SEC-MALS; Figure 1D-F). Together, these data suggest that the Red1 C-terminal domain forms coiled-coil homotetramers that associate end-to-end to form extended filaments.

The *S. cerevisiae* Red1 C-terminal domain was poorly-behaved *in vitro*, precluding an analysis of its oligomerization state and the effects of mutating residues I743 and I758. We instead mutated the equivalent residues in *Zr* Red1, I715 and M730, to arginine and examined the effects by SEC-MALS. We found that the *Zr* Red1^705-798^ I715R mutant formed a homotetramer in solution, instead of the extended filaments formed by wild-type *Zr* Red1^705-798^ (Figure 1D-E). The *Zr* Red1^705-798^ M730R mutant was poorly behaved in solution, precluding a detailed analysis of this mutant’s effects on filament formation.

The above data showing that mutation of *Zr* Red1 I715 disrupts filament assembly suggest that filament formation by Red1, rather than simply coiled-coil tetramer formation, may be critical for meiotic chromosome axis structure and function. To test this idea, we replaced the coiled-coil region of *S. cerevisiae* Red1 (residues 734-827) with the equivalent region of *Zr* Red1 (residues 707-798). We found that this chimeric Red1 protein localized to meiotic chromosomes and supported an overall spore viability of 54% (Figure 1H), significantly higher than prior measurements of spore viability for *redl△* strains (56, 57). Thus, the *Sc-Zr* Red1 chimera, while compromised relative to wild-type Red1, supports axis assembly, crossover formation, and meiotic chromosome segregation to a significant degree. We next removed the C-terminal seven residues of the chimeric Red1 to eliminate filament formation while maintaining Red1 homotetramers, and found that while this chimera localized to meiotic chromosomes, spore viability dropped to 2% (Figure 1H). These data support the idea that Red1 filament formation is critically important to the success of meiosis, and that the effects of the *Sc* Red1 I743A mutation, and likely also the I758R mutation, may be due to disruption of Red1 filament assembly.

### SYCP2 is an interaction hub for the mammalian chromosome axis

The mammalian chromosome axis comprises cohesin complexes with several meiosis-specific subunits (58-61); two meiotic HORMAD proteins, HORMAD1 and HORMAD2 (62, 63); and the coiled-coil proteins SYCP2 and SYCP3 (64-66). We have previously shown that both HORMAD1 and HORMAD2 possess closure motifs that associate with these proteins’ N-terminal HORMA domains (67), but no closure motifs on other axis proteins have been identified. SYCP2 has been proposed as a distant homolog of budding-yeast Red1, and possesses a similar domain structure: an N-terminal ordered domain that may mediate the protein’s association with chromosomes (68), followed by an extended disordered region and a C-terminal domain of ~175 residues predicted to form a coiled-coil. Instead of self-associating like Red1, however, the SYCP2 coiled-coil domain binds the shorter coiled-coil protein SYCP3 (Figure 2A) (50, 69, 70). Additionally, co-expression of SYCP2 and SYCP3 in cultured cells results in the assembly of large filamentous structures that incorporate both proteins, suggesting a capacity for self-assembly of SYCP2:SYCP3 complexes (31).

**Figure 2.**
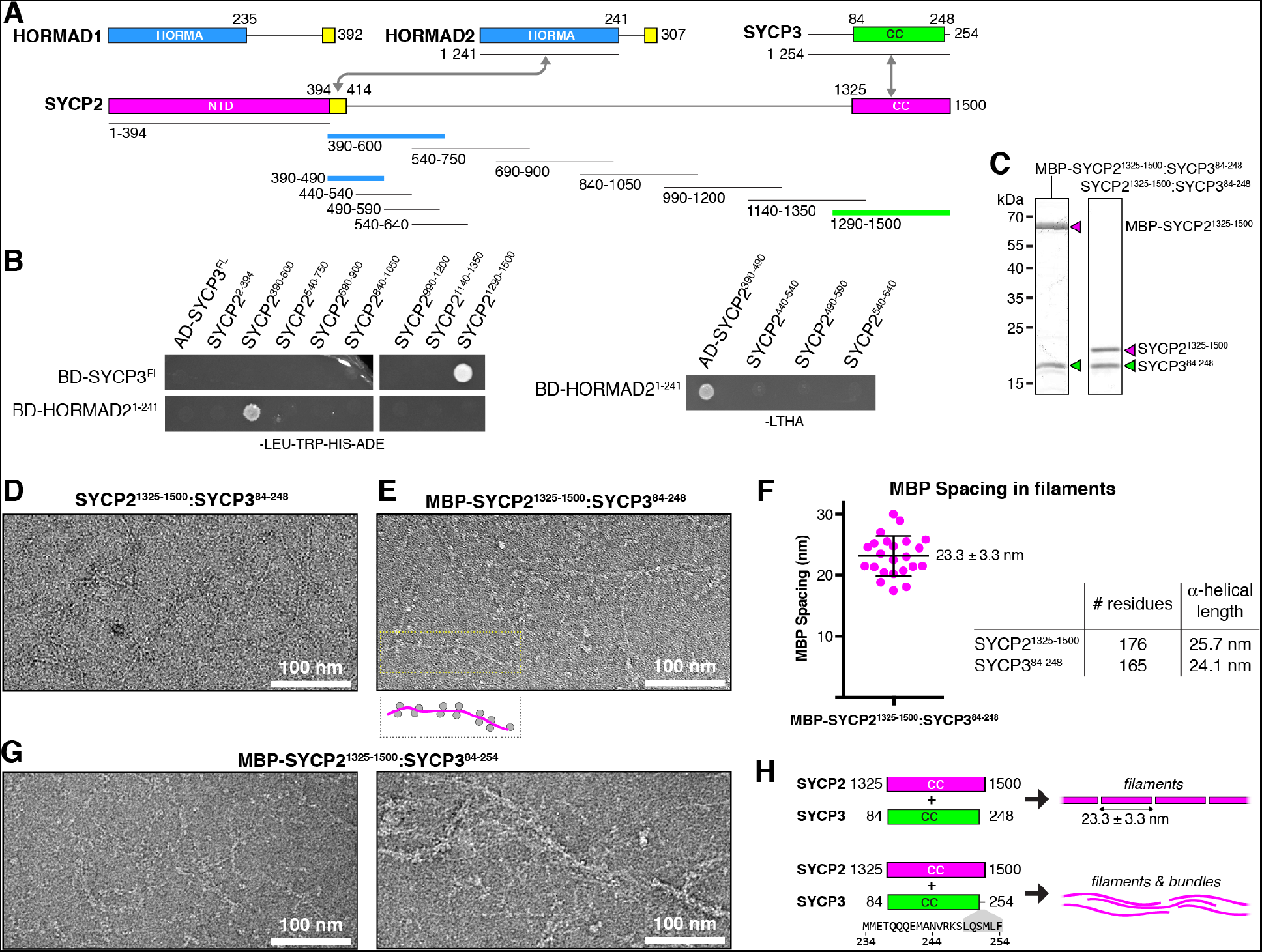
Protein-protein interactions and filament formation by mammalian SYCP2 and SYCP3. (A) Schematic ofM *musculus (Mm*) chromosome axis proteins, with underlines indicating fragments used for yeast two-hybrid analysis. The SYCP2 NTD (residues 1-394) forms a globular structure with unknown function (Feng et al. 2017). See Figure 2 - Figure Supplement 1 for detailed coiled-coil and alpha-helix predictions of the SYCP2 and SYCP3 C-terminal domains. Yellow regions indicate likely HORMAD-binding closure motifs. (B) Yeast two-hybrid analysis of SYCP2 truncations versus SYCP3 and the HORMAD2 HORMA domain. AD: Gal4 activation domain fusion; BD: Gal4 DNA-binding domain fusion. Stringent selection on -LEU-TRP-HIS-ADE (-LTHA) media is shown; see Figure 2 – Figure Supplement 2 for complete yeast two-hybrid results and for coexpression of SYCP2 fragments with HORMAD2^1-241^. (C) SDS-PAGE analysis of purified *Mm* SYCP2^1325-1500^:SYCP3^84-248^ complexes, with an N-terminal MBP tag on SYCP2 (left) or with the tag removed (right). (D) Representative negative-stain electron micrograph of purified *Mm* SYCP2^1325-1500^:SYCP3^84-248^. See Figure 2 – Figure Supplement 3A for additional full micrographs. (E) Representative negative-stain electron micrographs of purified His_6_-MBP-Mm SYCP2^1325-1500^:SYCP3^84-248^. See Figure 2 – Figure Supplement 3B for additional full micrographs. (F) Quantification of inter-MBP spacing in micrographs of His_6_-MBP-Mm SYCP2^1325-1500^:SYCP3^84-248^ filaments. The measured spacing of 23.1 +/- 3.3 nm is equivalent to the length of a ~160-residue coiled-coil (0.146 nm rise per residue). (G) Representative negative-stain electron micrographs of purified His_6_-MBP-Mm SYCP2^1325-1500^:SYCP3^84-254^. See Figure 2 – Figure Supplement 4 for additional full micrographs. *Right:* Quantification of inter-MBP spacing in micrographs of His_6_-MBP-Mm SYCP2^1325-1500^:SYCP3^84-248^ filaments. (H) Schematic summary of negative-stain electron microscopy results: *Mm* SYCP2^1325-1500^:SYCP3^84-248^ forms individual filaments assembled from ~23 nm units, while re-addition of the highly-conserved C-terminal six residues of SYCP3 (249-254; shown in gray below schematic) causes self-association/bundling of these filaments.

To outline protein-protein interactions within the mammalian chromosome axis, we used yeast two-hybrid assays to test for interactions between SYCP2, SYCP3, and HORMAD2. We identified a short region of SYCP2 directly following the protein’s ordered N-terminal domain (residues 395-414, Figure 2B, Figure 2 – Figure Supplement 2) that binds the HORMAD2 HORMA domain, showing that this region contains a closure motif. The location of the SYCP2 closure motif—directly C-terminal to the ordered N-terminal domain—is equivalent to the location of the budding-yeast Red1 closure motif, lending support to the idea that SYCP2 and Red1 are homologs.

Our yeast two-hybrid assays also confirmed that the coiled-coil regions of SYCP2 and SYCP3 associate (Figure 2B). To characterize the structure and oligomeric state of the SYCP2:SYCP3 complex, we next sought to purify this complex. We co-expressed the coiled-coil domains of *M. musculus* SYCP2 (residues 1325-1500) and SYCP3 (residues 84-248), and found that the proteins form a stoichiometric complex (Figure 2C) that, like the Red1 C-terminal domain, forms large assemblies in vitro as judged by size-exclusion chromatography (Figure 3A, B). Analysis of *Mm* SYCP2^1325-1500^:SYCP3^84-248^ assemblies by negative-stain electron microscopy revealed extended filaments much like those we observed for *Zr* Red1^705-798^ (Figure 2D, Figure 2 – Figure Supplement 3A). When we visualized the same complex with SYCP2 tagged at its N-terminus with MBP (*Mm* MBP-SYCP2^1325-1500^:SYCP3^84^’^248^) we observed filaments decorated with regularly-spaced pairs of densities ~5 nm in diameter, equivalent to the expected size of a single ~43-kDa MBP monomer (Figure 2E, Figure 2 – Figure Supplement 3B). We measured the inter-MBP spacing along SYCP2:SYCP3 filaments, and found an average spacing of 23.1 nm, which closely matches the expected length of an a-helical coiled coil ~175 residues in length (Figure 2F). These data suggest that the SYCP2:SYCP3 complex forms filaments through end-to-end association of individual a-helical units ~23 nm in length, with each unit containing two copies of SYCP2.

**Figure 3.**
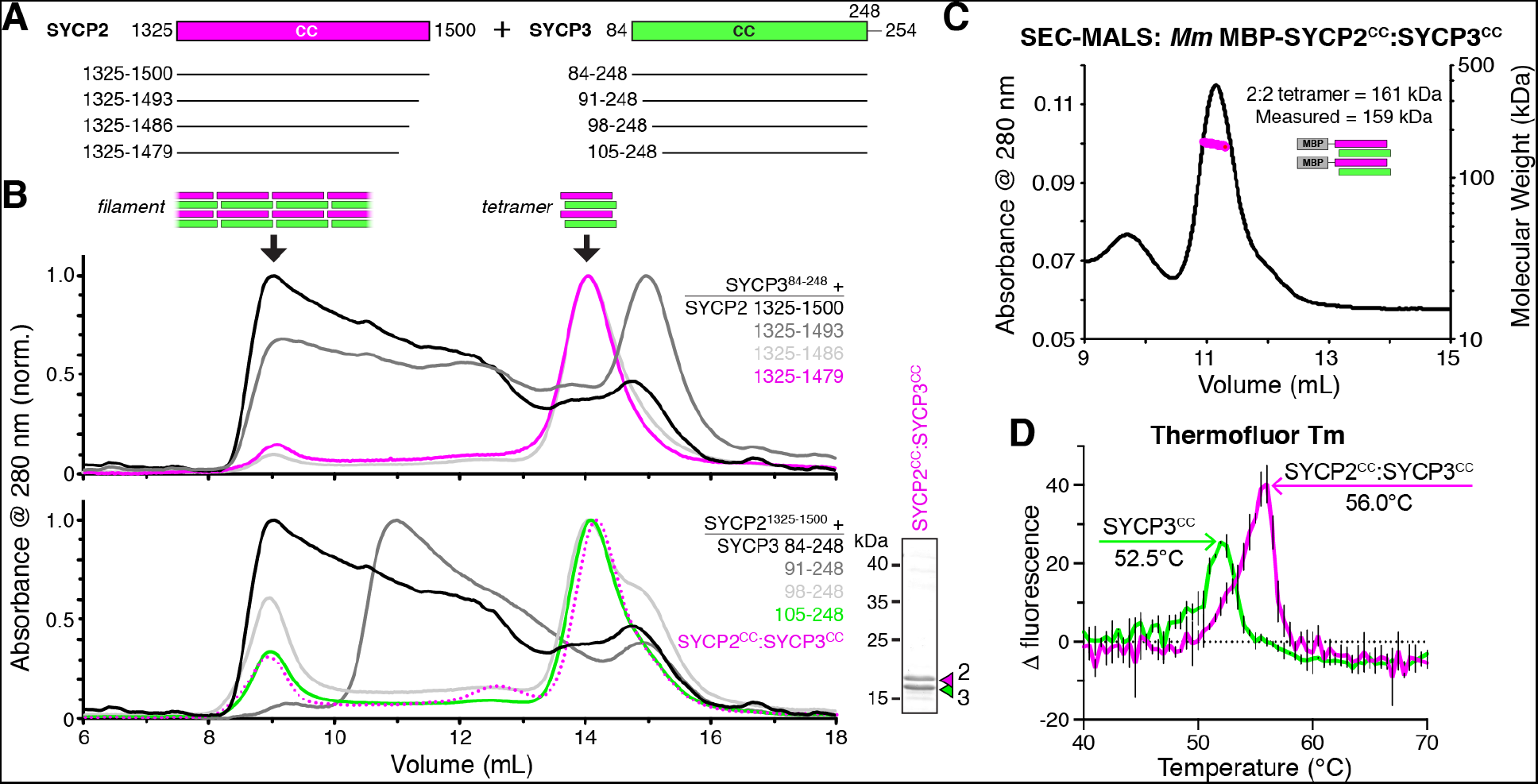
SYCP2:SYCP3 filaments are assembled from end-to-end interactions of a 2:2 heterotetrameric unit. (A) Schematic of the predicted coiled-coil regions of *M. musculus* SYCP2 and SYCP3, with truncations used for co-expression/size exclusion chromatography analysis in panels (B) and (C). (B) Superose-6 size exclusion chromatography analysis of truncated *Mm* SYCP2:SYCP3 complexes. All complexes were purified after co-expression using an N-terminal His_6_-MBP tag on SYCP2. *Upper panel:* truncation of the SYCP2 coiled-coil C-terminus, from 1325-1500 (black) to 1325-1479 (magenta), all co-expressed with SYCP3^84-248^. *Lower panel:* truncation of SYCP3 coiled-coil N-terminus, from 84-248 (black) to 105-248 (green), all co-expressed with SYCP31325-1500. Magenta dotted line indicates elution profile of MBP-SYCP2^1325-1479^:SYCP3^105-248^ complex (*Mm* SYCP2^cc^:SYCP3^cc^), used for SEC-MALS in panel (C). *Lower right:* SDS-PAGE analysis of purified *Mm* SYCP2^CC^:SYCP3^CC^ complex (with His_6_-MBP tag removed). (C) SEC-MALS analysis of purified His_6_-MBP-Mm SYCP2^cc^:SYCP3^cc^ complex. Calculated molecular weight of a 2:2 heterotetramer = 161.4 kDa; Measured molecular weight = 158.6 kDa. (D) Thermofluor melting-temperature (*Tm*) analysis for *Mm* SYCP2^cc^:SYCP3^cc^ (red) versus a homotetrameric *Mm* SYCP3^CC^ complex (green). Thick colored lines represent an average of three independent measurements, with standard deviation represented by thin vertical black lines.

The experiments above were conducted with a construct of SYCP3, residues 84-248, lacking the C-terminal six residues of this protein. These residues have been previously shown to be critical for formation of large homotypic SYCP3 filaments when overexpressed in mammalian tissue-culture cells (50, 71), and for formation of large SYCP3 assemblies in vitro (49). We next purified an SYCP2:SYCP3 complex containing these residues*,Mm* SYCP2^1325-1500^:SYCP3^84-254^, and visualized the complex by negative-stain electron microscopy. We found that this complex forms filaments equivalent to *Mm* SYCP2^1325-1500^:SYCP3^84^’^248^, but that in contrast to the truncated complex, filaments containing the full SYCP3 C-terminus tended to self-associate into bundles (Figure 2G). Given the high conservation of these residues and their importance for large-scale SYCP3 assembly in multiple assays, we propose that the SYCP3 C-terminus mediates bundling of SYCP2:SYCP3 filaments as a critical step in assembly of the meiotic chromosome axis (Figure 2H).

We next sought to further dissect the SYCP2:SYCP3 filament assembly. We progressively truncated both proteins, and found that removal of 21 residues from either the SYCP2 C-terminus (residues 1480-1500) or the SYCP3 N-terminus (residues 84-104) resulted in a complete loss of large assemblies, while maintaining the association between the two proteins (Figure 3A,B). We combined the truncations on both proteins to yield a minimal construct, *Mm* SYCP2^1325-1479^:SYCP3^105-248^, which we term SYCP2^cc^:SYCP3^cc^ hereon. We first measured the molecular weight of the SYCP2^CC^:SYCP3^CC^ complex with an N-terminal MBP tag on SYCP2^CC^ by SEC-MALS. The measured molecular weight of this complex, 159 kDa, closely matched the predicted molecular weight of 161 kDa for a 2:2 heterotetramer of SYCP2 and SYCP3 (Figure 3C).

Prior work on *H. sapiens* SYCP3 has shown that this protein self-associates to form coiled-coil homoetramers in vitro (49). We purified *Mm* SYCP3^CC^ in the absence of SYCP2 and determined its structure by x-ray crystallography, revealing an antiparallel coiled-coil homotetramer similar in structure to *H. sapiens* SYCP3 (Figure 4 – Figure Supplement 1). As SYCP2 and SYCP3 share limited sequence homology in their coiled-coil region, we reasoned that SYCP3 homotetramers may form through promiscuous coiled-coil interactions in the absence of SYCP2. To compare the stability of *Mm* SYCP3^CC^ homotetramers with *Mm* SYCP2^CC^:SYCP3^CC^ heterotetramers, we measured their melting temperatures (Tm) using a dye-binding assay. We found that the SYCP2^CC^:SYCP3^CC^ heterotetramer is more stable than SYCP3^CC^ on its own (56.0°C *Tm* versus 52.5°C; Figure 3D), supporting the idea that the heterotetrameric complex is the preferred state when both proteins are present.

**Figure 4.**
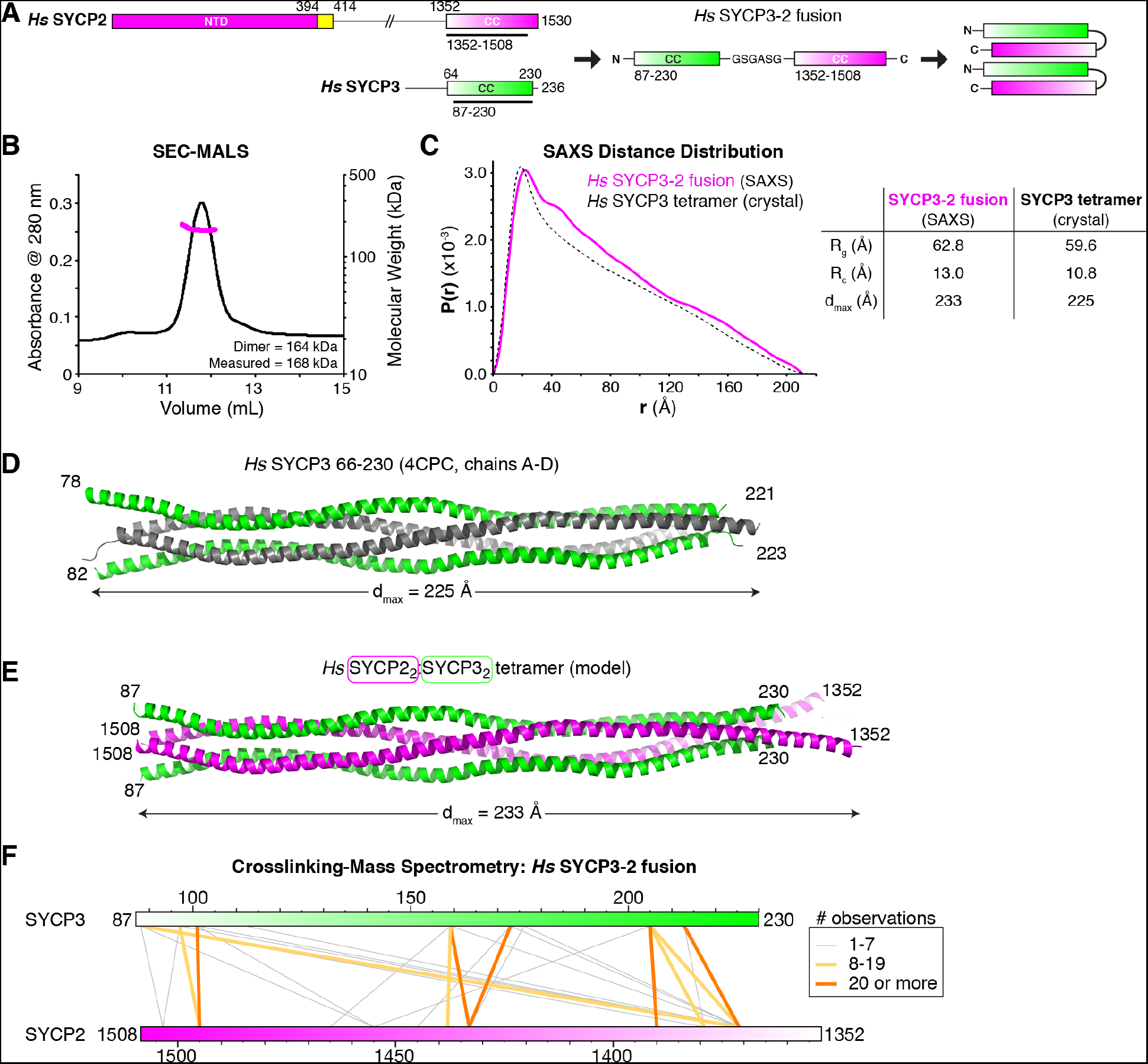
Structural analysis of the SYCP2:SYCP3 complex. (A) Design of the *H. sapiens* SYCP3^cc^-SYCP2^cc^ fusion, based on the idea that SYCP2 and SYCP3 helices pack antiparallel in the 2:2 heterotetrameric structure. (B) SEC-MALS analysis of purified His_6_-MBP-Hs SYCP3^CC^-[GSGASG]-SYCP2^CC^. Measured molecular weight (167.7 kDa) is equivalent to the calculated molecular weight of a homodimer (equivalent to a 2:2 heterotetramer of SYCP2 and SYCP3; 163.7 kDa). (C) Intra-particle distance distribution (*P(r)*) curve for the *Hs* SYCP3^CC^-SYCP2^CC^ fusion construct derived from small-angle x-ray scattering (SAXS) analysis (magenta), compared to the calculated distance distribution of the *Hs* SYCP3^CC^ homotetramer structure (PDB ID 4CPC; dotted black line) (49). Lower: table comparing radius of gyration (*Rg*), cross-sectional radius of gyration (Rc), and maximum dimensions (*dmax*) of the *Hs* SYCP3^CC^-SYCP2^CC^ fusion (calculated from SAXS; see Figure 4 – Figure Supplement 2) and the *Hs* SYCP3^CC^ homotetramer (calculated from the crystal structure). See Figure 4 – Figure Supplement 3 for SAXS analysis of the *Mm* SYCP2^CC^:SYCP3^CC^ complex, which showed similar results but was affected by aggregation. (D) Structure of the *Hs* SYCP3^CC^ homotetramer structure (PDB ID 4CPC; dotted black line) (49), with two parallel chains (N-termini left) colored green, and the other two chains (N-termini right) colored gray. We determined the crystal structure of the*M. musculus* SYCP3CC homotetramer in two different crystal forms (Figure 4 – Figure Supplement 1A-D). This structure resembles the structure of *Hs* SYCP3^CC^ in the central coiled-coil region, but adopts a distinct, more disordered structure near both ends. (E) Model of an *Hs* SYCP2^CC^:SYCP3^CC^ 2:2 heterotetramer, with two SYCP3 chains colored green as in panel (C) (N-termini left), and two SYCP2 chains colored magenta (N-termini right). Sequence register was derived from aligning SYCP2 and SYCP3 sequences (schematized in Figure 4 – Figure Supplement 1E). (F) Schematic of crosslinking mass spectrometry (XLMS) results on the *Hs* SYCP3^CC^-SYCP2^CC^ fusion. Crosslinks observed at least 8 times are colored yellow, and crosslinks observed at least 20 times are colored orange. See Tables S2 and S3, and Figure 4 – Figure Supplement 4 for full results. 9 of 10 high-scoring crosslinks support the antiparallel subunit arrangement shown in panel (D).

### The SYCP2:SYCP3 complex is an antiparallel heterotetramer

While the SYCP3^CC^ homotetramer is likely not the favored state in the presence of SYCP2, its structure may nonetheless be informative as to the structure of SYCP2^CC^:SYCP3^CC^. Given its 2:2 stoichiometry and our observed effects on filament formation from truncating opposite ends of SYCP2 and SYCP3, we reasoned that SYCP2^CC^:SYCP3^CC^ may form a complex with two SYCP2 protomers oriented antiparallel to two SYCP3 protomers. To test this idea, we generated a series of *Hs* and *Mm* SYCP2:SYCP3 constructs with the two proteins fused end-to-end through a short peptide linker (Figure 4A). One such construct, *Hs* SYCP3^87^’^230^-[GSGASG]-SYCP2^1352^’ ^1508^ (termed *Hs* SYCP3^CC^-SYCP2^CC^ fusion hereon), was highly-expressed in *E. coli* and formed a stable dimer by SEC-MALS, equivalent to an SYCP2:SYCP3 heterotetramer (Figure 4B). We were unable to crystallize this complex, so we turned instead turned to small-angle x-ray scattering, which provides low-resolution size and shape information on macromolecular complexes in solution. SAXS can provide a reliable measure of a particle’s maximum dimension (*dmax*) and radius of gyration (*Rg)*, as well as, for cylindrical particles, the cross-sectional radius of gyration (Rc) (72, 73). Analysis of the *Hs* SYCP3^CC^-SYCP2^CC^ fusion by SAXS showed that this complex’s *dmax, Rg*, and *Rc* closely match theoretical values calculated from the crystal structure of the *Hs* SYCP3^CC^ homotetramer (Figure 4C-E). Further, the intra-particle distance distribution function calculated from the SAXS scattering curve also closely matched the profile calculated from the *Hs* SYCP3^CC^ crystal structure (Figure 4C). We also performed SAXS analysis on the heterotetrameric *Mm* SYCP2^CC^:SYCP3^CC^ complex (Figure 4 – Figure supplement 3). This complex partially aggregated in solution, precluding detailed analysis, but showed results broadly consistent with the *Hs* SYCP3^cc^-SYCP2^cc^ fusion. Overall, these data show that the SYCP2^CC^:SYCP3^CC^ complex forms an extended coiled-coil tetramer with an overall structure similar to that of the SYCP3 homotetramer.

To test the idea that SYCP2 and SYCP3 are oriented anti-parallel to one another in the SYCP2^CC^:SYCP3^CC^ complex, we used cross-linking mass spectrometry (XLMS), which identifies pairs of lysine residues whose side-chains are in close proximity in a native complex. We identified 55 cross-links in the *Hs* SYCP3^CC^-SYCP2^CC^ fusion construct: 15 within the SYCP3 region, 13 within SYCP2, and 27 between SYCP3 and SYCP2 (Table S2, S3). Of the 27 cross-links identified between SYCP2 and SYCP3, ten were observed at least 8 times in our mass spectrometry experiments. When mapped onto either parallel or antiparallel models of SYCP2:SYCP3, nine of these ten crosslinks supported an antiparallel arrangement of SYCP2 and SYCP3 α-helices (Figure 4F, Figure 4 – Figure Supplement 4). When examining intra-SYCP3 and intra-SYCP2 crosslinks, the majority of robustly-observed crosslinks were between residues close in sequence, as expected if the two protomers of each protein are oriented parallel to one another (Figure 4 – Figure Supplement 4). When combined with the above SAXS data, these data strongly support a model in which SYCP2 and SYCP3 form an antiparallel, heterotetrameric coiled-coil complex. We propose that heterotetrameric SYCP2:SYCP3 complexes associate end-to-end to form extended filaments, which can then bundle through the SYCP3 C-terminus to form the chromosome axis.

### Plant ASY3 binds HORMADs and forms filaments with ASY4

In higher plants, the chromosome axis comprises meiosis-specific cohesin complexes (74, 77); two meiotic HORMAD proteins, ASY1 and ASY2 (24, 78): and two coiled-coil proteins, ASY3 and ASY4 (33, 34). ASY3 is required for axis localization of ASY1, and its disruption causes a strong defect in crossover formation (33). Despite low sequence identity with either Red1 or SYCP2, ASY3 has been proposed as a functional homolog of Red1 based on phenotypic similarities plus the presence of a conserved C-terminal domain with predicted coiled-coil character (Figure 5A) (33). ASY4 was recently identified by two groups as a short protein with high homology to the ASY3 coiled-coil domain, that also interacts with ASY3 (34, 79).

**Figure 5.**
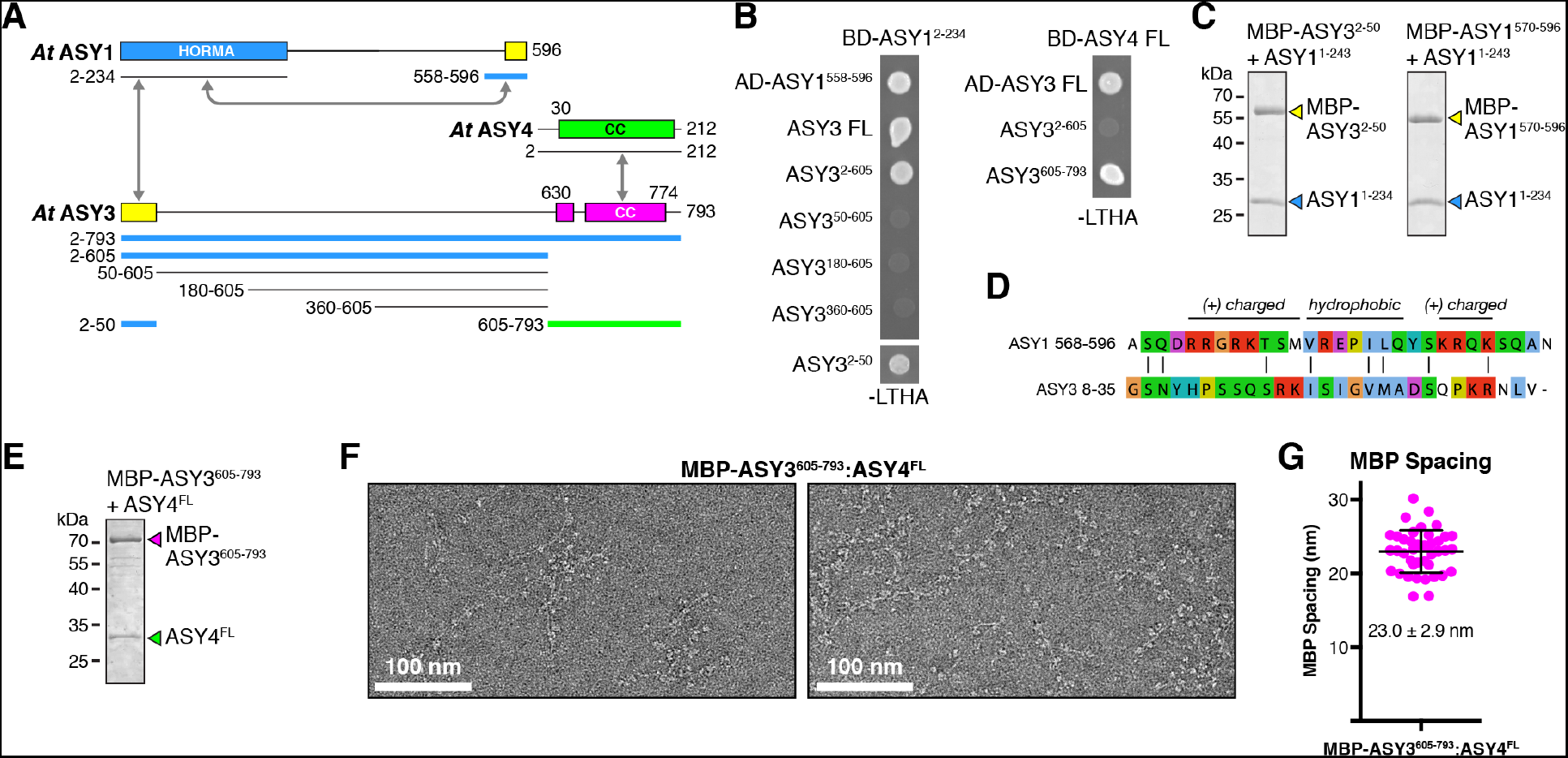
Plant ASY3 and ASY4 share a conserved molecular architecture with mammalian and budding-yeast axis proteins. (A) Schematic of *Arabidopsis thaliana* chromosome axis proteins, with truncations used for yeast two-hybrid assays shown for ASY1, ASY3, and ASY4. Colored in blue and green are ASY3 constructs that interact with the ASY1 HORMA domain (blue) and ASY4 (green). (B) Yeast two-hybrid analysis of *A. thaliana* ASY1, ASY3, and ASY4. AD: Gal4 activation domain fusion; BD: Gal4 DNA-binding domain fusion. Stringent selection on -LEU-TRP-HIS-ADE (-LTHA) media is shown; see Figure 5 – Figure Supplement 1 for complete results. (C) SDS-PAGE analysis of purified His_6_-MBP-tagged closure motifs in ASY3 (residues 2-50) and ASY1 (residues 570-596) in complex with untagged ASY1 HORMA domain (residues 1-234). Complexes were purified using Ni^2+^ affinity and size-exclusion chromatography. (D) Sequence alignment of the putative closure motif regions of *At* ASY1 (residues 568-596) and ASY3 (residues 8-35). The two regions show weak homology with a central region enriched in hydrophobic residues, bracketed on both sides by positively-charged residues. See Figure 5 – Figure Supplement 2 for sequence alignments of both regions. (E) SDS-PAGE analysis of purified His_6_-MBP-ASY3^605-793^:ASY4^FL^ complexes used for negative-stain EM analysis (panel F). (F) Representative negative-stain electron micrographs of purified His_6_-MBP-ASY3^605-793^:ASY4^FL^ filaments. See Figure 5 – Figure Supplement 3 for additional full micrographs. (G) Quantification of inter-MBP spacing in micrographs of His_6_-MBP-Mm SYCP2^1325-1500^:SYCP3^84-248^ filaments. The measured spacing of 23.0 +/- 2.9 nm is equivalent to the length of a ~160-residue coiled-coil (0.146 nm rise per residue). Predicted coiled-coil regions of ASY3 and ASY4 are ~145 and ~180 residues, respectively.

To define protein-protein interactions within the plant chromosome axis, we used yeast two-hybrid assays to test interactions between *A. thaliana* ASY1, ASY3, and ASY4. We found that the ASY1 N-terminal HORMA domain (residues 1-234) interacts with its own extreme C-terminus (residues 558-596), revealing that this protein possesses a C-terminal closure motif like its orthologs in *C. elegans*, mammals, and fungi (Figure 5B). We further identified an ASY1 HORMA domain-interacting region at the N-terminus of ASY3 (residues 1-50; Figure 5B). This region contains a highly-conserved motif of ~30 residues with limited sequence homology to the ASY1 C-terminus (Figure 5D, Figure 5 – Figure Supplement 2), suggesting that both regions act as HORMAD-binding closure motifs. To verify these interactions, we co-expressed each putative closure motif (fused to an N-terminal His_6_-MBP tag) with the ASY1 HORMA domain in *E. coli*. Both His_6_-MBP-ASY3^2-50^ and His_6_-MBP-ASY1^570-596^ co-purified with untagged ASY1 HORMA domain through Ni^2+^-affìnity and size exclusion chromatography (Figure 5C), demonstrating a high-affinity interaction. These findings show that plant meiotic HORMADs, like those from fungi and mammals, can interact with closure motif sequences both at their own C-termini and in the N-terminal region of a Red1-like axis core protein.

We next tested interactions between *A. thaliana* ASY3 and ASY4. We found that the C-terminal coiled-coil region of *At* ASY3 (residues 605-793) interacts with ASY4, confirming the recent finding of Osman et al. in *Brassica oleracea* (34) and of Chambon et al. in *A. thaliana* (79) (Figure 5B). We next purified a complex between the coiled-coil domains of *A. thaliana* ASY3 and ASY4 (Figure 5E), which formed large assemblies in solution as measured by size-exclusion chromatography. Negative-stain electron microscopy on purified His_6_-MBP-ASY3^605-793^:ASY4^FL^ assemblies revealed extended filaments equivalent to those observed with both budding-yeast Red1 and mammalian SYCP2:SYCP3 (Figure 5F). As with the *Mm* MBP-SYCP2^1325^’^1500^:SYCP3^84^’^248^ filament, these filaments were decorated at regular intervals with pairs of MBP densities. When we measured the average distance between paired MBP densities along these filaments, we obtained an average spacing of 23.0 +/- 2.9 nm, in close agreement with the spacing in SYCP2:SYCP3 filaments and with the predicted length of the ASY3 and ASY4 coiled-coil regions (~145 and ~180 residues, respectively, corresponding to coiled-coil lengths of ~21.2 and ~26.3 nm; Figure 5G). These data strongly suggest that ASY3 and ASY4 assemble into 2:2 heterotetramers that associate end-to-end to form extended filaments, in a manner equivalent to both mammalian SYCP2:SYCP3 and fungal Red1.

## Discussion

The meiotic chromosome axis plays several crucial roles to support inter-homolog crossover formation and signaling in meiosis I. The first major role is to provide a scaffold for organization of each chromosome as a linear array of loops, with these loops directly extruded or otherwise constrained by cohesin complexes (1). The axis assembles in early meiotic prophase, when chromosomes just become visible as the “thin threads” for which the leptotene stage is named. As cells progress through zygotene and then pachytene (“thick threads”), chromosomes undergo significant linear compaction without disruption of the underlying chromosome axis structure. We have shown here that budding-yeast Red1, mammalian SYCP2:SYCP3, and plant ASY3:ASY4 all form filaments from homo- or hetero-tetrameric units, and that the SYCP2:SYCP3 filaments have a tendency to form bundles. We propose that individual short filaments of these “axis core proteins” associate loosely with cohesin complexes, then form bundles to assemble a flexible scaffold for cohesin-mediated extrusion/constraint of chromatin loops (Figure 6). In this scheme, both filament formation by axis core proteins and cohesin activity are required for axis assembly and chromosome compaction, explaining how mutation of axis core proteins like SYCP3 (28, 29, 80) or cohesin subunits including SMC1β (80, 81), REC8 (82, 83), RAD21L (59, 84), and STAG3 (59-61, 85-87) can affect the overall length of the axis. An axis constructed from a flexible core of loosely-associated filaments would also enable axis extension or compression in processes like synaptic adjustment, in which the lengths of two chromosomes can adjust to one another as the synaptonemal complex (SC) assembles between them (1, 88).

**Figure 6.**
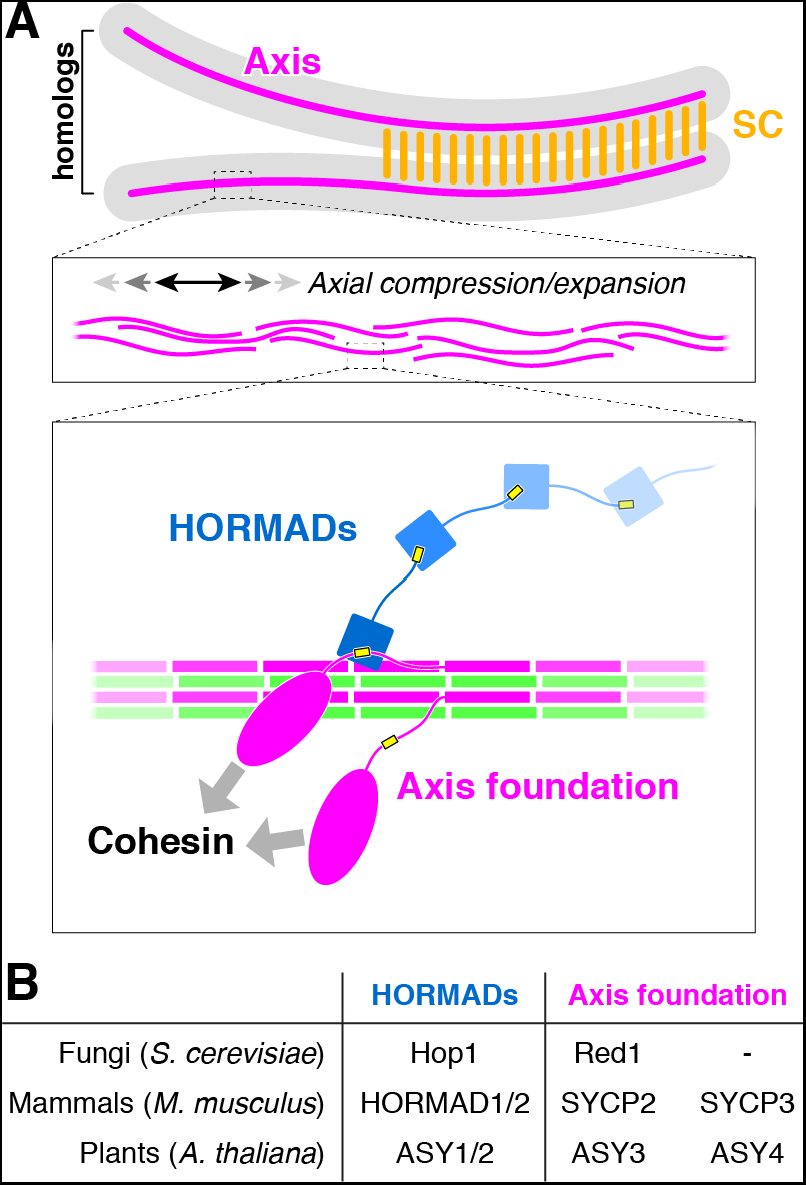
The conserved molecular architecture of the meiotic chromosome axis. (A) Model for assembly of the meiotic chromosome axis in fungi, plants, and mammals. Related axis core proteins (fungal Red1, plant ASY3:ASY4, mammalian SYCP2:SYCP3) form filaments from coiled-coil homo- or heterotetrameric units, which flexibly associate with chromosome-associated cohesin complexes. Chromatin loop extrusion by cohesin complexes and bundling of axis-core filaments leads to formation of the chromosome axis, which is flexible and able to axially compress or expand if needed. (B) List of homologous chromosome axis proteins in different eukaryotic groups.

A second critical function of the chromosome axis is to promote the formation of meiotic DSBs, then orchestrate the repair of a subset of DSBs as inter-homolog crossovers. In most organisms, the meiotic HORMADs are major regulators of both DSB formation and crossover formation. We have previously proposed an overall axis assembly pathway in *S. cerevisiae* with cohesin-associated Red1 recruiting Hop1 through its closure motif (25), and we can now extend this model to both mammals and plants. We propose that a key conserved function of the axis core proteins is to recruit meiotic HORMADs through their HORMA domain-binding closure motifs. These localized HORMADs may then recruit additional HORMADs through head-to-tail oligomer formation through their own C-terminal closure motifs. As cells enter pachytene and the axis core proteins become integrated into the SC, HORMADs are removed from the axis by the Pch2/TRIP13 ATPase, thereby suppressing further DSB formation and licensing the progression of meiosis (19, 36-39).

The third major function of the meiotic axis is to serve as the lateral elements of the SC in pachytene, after the bulk of meiotic HORMADs have been removed. Our physical model of the mammalian chromosome axis, comprising a bundle of SYCP2:SYCP3 filaments with a periodicity of 23 nm, generally agrees with prior electron microscopy (EM) studies showing ~20 nm periodicity in the lateral elements of assembled SCs (89). Also in agreement with this model, a recent analysis of mouse SC structure by super-resolution light microscopy has shown that SYCP3 and the SYCP2 coiled-coil region perfectly co-localize in a single “cable” in each lateral element, and that the C-terminus of the transverse filament protein SYCP1 is situated close to this cable (48). Significant questions remain regarding how the SC lateral elements and transverse filaments might interact, and the role of cohesin complexes in this interaction is also unknown. Recently, it was reported that REC8 and RAD21L, meiosis-specific cohesin complex kleisin subunits, both localize slightly “inside” SYCP3; that is, they are situated closer to the SYCP1 transverse filaments than the SYCP2:SYCP3 complex (90). These data suggest that cohesin complexes may somehow be integrated into the structure of the SC, in a manner that is not yet well-understood.

Several recent studies have reported that in vitro, mammalian SYCP3 forms coiled-coil homotetramers that further assemble into large oligomeric structures with a 22 nm periodicity (49, 51, 52). Further, SYCP3 can bind DNA through two short patches of basic residues near the N-terminus of this protein’s coiled-coil domain (49), and large SYCP3 oligomers appear to bind and condense plasmid DNA (52). These data have led to a model whereby homotypic SYCP3 oligomers, interacting directly with DNA, form a major part of the mammalian chromosome axis (49, 52). While we cannot rule out the formation of homotypic SYCP3 assemblies in meiotic cells, our data shows that the SYCP2:SYCP3 heterotetramer is more stable in solution than the SYCP3 homotetramer, and is therefore likely to be the preferred assembly in meiotic cells. Further, as SYCP3 does not localize to the chromosome axis in a mutant of SYCP2 lacking its coiled-coil domain (30), direct SYCP3-DNA binding is unlikely to contribute significantly to axis formation.

We have shown that the architecture of the meiotic chromosome axis is highly conserved across fungi, mammals, and plants. Our model assigns critical functions in both overall axis structure and HORMAD recruitment to the axis core proteins, yet some organisms, including *C. elegans* and *D. melanogaster*, appear to lack axis core proteins entirely. Our prior work has strongly suggested that the meiotic HORMADs in *C. elegans* interact directly with cohesin complexes (67), and thus far meiotic HORMADs have not been identified in *D. melanogaster*. We propose that the key feature of meiosis in both *C. elegans* and *D. melanogaster* that eliminates the need for axis core proteins is that these organisms assemble the SC prior to meiotic recombination. Thus, the SC itself can provide a physical scaffold for chromosome organization and recombination control in these organisms, eliminating much of the need for a distinct chromosome axis that assembles prior to SC formation.

While our data shed significant new light on the assembly and function of the meiotic chromosome axis, significant questions remain. First, while the N-terminal domains of both Red1 and SYCP2 likely adopt similar structures and mediate these proteins’ association with meiotic chromosomes, their direct binding partners are as yet mysterious. A recent study identified several potential binding partners of the SYCP2 N-terminal domain (68), but as these proteins are mostly centromere-associated, it remains unknown what SYCP2 may bind along the length of chromosomes. We propose that the SYCP2 and Red1 N-terminal domains may bind cohesin complexes directly (91), or may instead bind one or more chromatin-associated proteins, perhaps even a specific histone mark, to mediate a flexible interaction with chromatin.

A further mystery involves plant ASY3, which appears to entirely lack a Red1/SYCP2-like N-terminal domain. This protein may associate with chromosome-localized proteins through one or more short conserved motifs in its extended disordered region, or may in fact be recruited through interactions with the HORMADs ASY1 and ASY2. Both plant ASY1 and fungal Hop1 possess putative DNA- or protein-binding domains in their central regions, raising the possibility that these meiotic HORMADs could recruit axis core proteins to meiotic chromosomes, in a reversal of the canonical localization-dependence of these proteins. Thus, while the overall theme of axis assembly through filament formation is likely conserved across many eukaryotic families, each family appears to have evolved variations on this theme in keeping with its own unique requirements.

## Materials and Methods

### Cloning and Protein Purification

#### Mammalian proteins

For yeast two-hybrid analysis, *M. musculus* genes were PCR-amplified from cDNA (SYCP2: Harvard PlasmID clone MmCD00083242; HORMAD1: TransOMIC technologies clone BC051129; HORMAD2: TransOMIC technologies clone BC120781) or synthesized DNA fragment (SYCP3; GeneArt) and inserted by ligation-independent cloning into modified pBridge and pGADT7 vectors (Clontech). For co-expression, *M. musculus* SYCP2 and SYCP3 fragments were separately cloned into UC Berkeley Macrolab vectors 2CT (SYCP2; Amp^R^, N-terminal His_6_-MBP fusion) or 13S-A (SYCP3; Spec^R^, no tag) by ligation-independent cloning. For expression of SYCP3 alone, *M. musculus* SYCP3^105-248^ was cloned into UC Berkeley Macrolab vector 2ST (N-terminal His_6_-SUMO fusion). The *H. sapiens* SYCP2-SYCP3 fusion construct was assembled by multi-part PCR from synthesized-fragment templates (GeneArt) and inserted into vector 2CT by ligation-independent cloning. For co-expression of *M. musculus* SYCP2:HORMAD2 complexes, a polycistronic expression cassette was assembled by PCR and inserted into vector 2CT, yielding a final vector encoding His_6_-MBP-tagged SYCP2 fragments plus untagged HORMAD2^1^’^241^.

For purification of SYCP2:SYCP3 complexes, His_6_-MBP-SYCP2 and SYCP3 constructs were co-transformed into *E. coli* strain Rosetta 2(DE3) pLysS (Novagen), and grown in the presence of ampicillin, spectinomycin and chloramphenical to an OD600 of 0.9 at 37°C, induced with 0.25 mM IPTG, then grown for a further 16 hours at 18°C prior to harvesting by centrifugation. For purification, cells were lysed by sonication, then clarified lysates were purified by Ni^2^+ affinity (HisTrap HP; GE Life Sciences), ion exchange (HiTrap SP or Q; GE Life Sciences), and size exclusion chromatography (Superdex 200; GE Life Sciences). *H. sapiens* SYCP2:HORMAD2 complexes were coexpressed as above, then purified by Ni^2^+ affinity chromatography and analyzed by SDS-PAGE. In cases where cleavage of N-terminal His_6_-MBP or His_6_-SUMO tags was required, tags were removed by incubation with TEV protease at 4°C for 16 hours, then the mixture was passed over a Ni^2+^ affinity column, and the flow-through fractions were concentrated and purified by size-exclusion chromatography.

For size-exclusion chromatography-based assays of SYCP2:SYCP3 filament formation, His_6_-MBP-SYCP2:SYCP3 complexes were initially purified by Ni-NTA chromatography, then passed over a Superose-6 size-exclusion column (GE Life Sciences) to remove small-molecular weight contaminants. Fractions corresponding to the entire range containing SYCP2:SYCP3 complexes were pooled, concentrated, then passed over Superose-6 a second time for the traces shown in Figure 3B. N-terminal His_6_-MBP tags were not removed for this analysis.

For size exclusion chromatography coupled to multi-angle light scattering (SEC-MALS), 100 μL purified proteins at 2-5 mg/mL was injected onto a Superdex 200 Increase 10/300 GL column (GE Life Sciences) in a buffer containing 20 mM HEPES pH 7.5, 300 mM NaCl, 5% glycerol, and 1 mM DTT. Light scattering and refractive index profiles were collected by miniDAWN TREOS and Optilab T-rEX detectors (Wyatt Technology), respectively, and molecular weight was calculated using ASTRA v. 6 software (Wyatt Technology).

#### Fungal proteins

*Zygosaccharomyces rouxii* Red1 constructs were amplified by PCR and inserted by ligation-independent cloning into UC Berkeley Macrolab vector 2CT (Amp^R^, N-terminal His_6_-MBP fusion) for expression in *E. coli*. Proteins were expressed and purified as above.

#### Plant proteins

Full-length codon-optimized genes for *Arabidopsis thaliana* ASY1, ASY3, and ASY4 were synthesized (GeneArt) and inserted by ligation-independent cloning into modified pBridge and pGADT7 vectors (Clontech) for yeast two-hybrid analysis, or cloned into UC Berkeley Macrolab vectors 2CT /13S-A for expression in *E. coli*. Truncations were amplified by PCR and similarly cloned.

For co-purification of the ASY1 HORMA domain with putative closure motif peptides, putative closure motifs (ASY3^2-50^ and ASY1^570-596^) in vector 2CT (N-terminal His_6_-MBP fusion) and ASY1^1-234^ in vector 13S-A (untagged) were co-transformed into *E. coli* strain Rosetta 2(DE3) pLysS, and grown in the presence of ampicillin, spectinomycin and chloramphenical to an OD600 of 0.9 at 37°C, induced with 0.25 mM IPTG, then grown for a further 16 hours at 18°C prior to harvesting by centrifugation. For purification, cells were lysed by sonication, then clarified lysates were purified by Ni^2+^ affinity (HisTrap HP; GE Life Sciences) and size exclusion chromatography (Superdex 200; GE Life Sciences). For purification of ASY3:ASY4 for electron microscopy, ASY3^605-793^ in vector 2CT (N-terminal His_6_-MBP fusion) and full-length ASY4 in vector 13S-A (untagged) were co-transformed into *E. coli* strain Rosetta 2(DE3) pLysS, and grown in the presence of ampicillin, spectinomycin and chloramphenical to an OD600 of 0.9 at 37°C, induced with 0.25 mM IPTG, then grown for a further 16 hours at 18°C prior to harvesting by centrifugation, and purified as above.

### Yeast two-hybrid

For yeast two-hybrid analysis, plasmids were transformed into AH109 and Y187 yeast strains (Clontech), and transformants were selected using CSM -Leu (for pGADT7 vectors) and CSM -Trp (pBridge vectors) media. Haploid yeast strains were mated overnight at room temperature, and diploids were selected using CSM -Leu-Trp media. Diploids were patched onto low-stringency (CSM -Leu-Trp-His) and high stringency media (CSM -Trp-Leu-His-Ade), grown for 1-3 days at 30°C, and imaged.

### Electron microscopy

For negative-stain electron microscopy, protein complexes were passed over a size exclusion column (Superdex 200 Increase 10/300 GL; GE Life Sciences) in EM buffer (300 mM NaCl, 20 mM Tris-HCl pH 7.5, 1 mM DTT), and peak fractions were diluted to ~0.01 mg/mL in EM buffer. Samples were spotted on freshly glow-discharged carbon coated copper grids, blotted into a thin film, and stained using 2% of uranyl formate. Electron micrographs were acquired on a Tecnai F20 Twin transmission electron microscope (FEI, Hillsboro OR) operating at 200 kV on a Tietz F416 4K x 4K CMOS camera (TVIPS, Gauting, Germany). For untagged *Zr* Red1^705-798^ and MBP-ASY3^605-793^:ASY4^FL^, micrographs were acquired on a FEI Talos F200C with 4K x 4K CMOS camera (Thermo Fisher Scientific). Micrographs of His_6_-MBP-SYCP2^1325-1500^:SYCP3^84-248^ and MBP-ASY3^605-793^:ASY4^FL^ were analyzed using ImageJ to determine the average spacing of MBP densities on the respective filaments.

### Thermofluor melting assays

For measurement of melting temperature, 45 uL 0.1 mg/mL purified protein in gel-filtration buffer (20 mM Tris-HCl pH 7.5, 300 mM NaCl, 10% glycerol, 1 mM DTT) was mixed with 5 uL 50X SYPRO orange dye (Life Technologies; 5X final concentration) and pipetted into an optically-clear qPCR plate. SYPRO fluorescence was measured in a Bio-Rad CFX96 qPCR machine in FRET mode (excitation 450-490 nm, emission 560-580) using a temperature range 25-95°C in 0.5° steps (15 second hold per step). Triplicate measurements were averaged, buffer-subtracted, then the derivative of the fluorescence was calculated. The maximum value of the derivative curve (highest rate of change in fluorescence) is assigned as the *Tm*. N-terminal His_6_-MBP and His_6_-SUMO on SYCP2^1325^’^1479^:SYCP3^105^’^248^ and SYCP3^105^’^248^, respectively, were removed prior to Tm analysis.

### Crystallization and structure determination of *M. musculus* SYCP3 homotetramer

When co-expressed in *E. coli, M. musculus* SYCP3 is expressed at much higher levels than SYCP2 (not shown). We found that while *M. musculus* SYCP2^CC^ is insoluble when expressed without SYCP3^CC^, SYCP3^CC^ is able to form soluble homotetramers. While optimizing expression constructs, we co-expressed *M. musculus* His_6_-SUMO-SYCP3^105-248^ with untagged SYCP2^1325-1472^, purified the resulting complex, and identified crystallization conditions. Crystals were obtained in hanging drop format by mixing protein (50-80 mg/mL) with two parts well solution containing 100 mM Tris-HCl pH 8.5, 16% PEG 4000, and 100-200 mM sodium acetate. Later analysis showed that these crystals contain SYCP3 homotetrameric complexes, rather than SYCP2:SYCP3 heterotetramers. Because of the tendency of SYCP3^CC^ to form homotetrameric complexes, all other analysis with SYCP2^CC^:SYCP3^CC^ complexes was performed with complexes expressed with tagged SYCP2 and untagged SYCP3.

SYCP3 homotetramer crystals were cryoprotected by the addition of 20% sucrose, then diffraction data was collected at the Advanced Photon Source, beamline 24ID-C. Despite identical growth conditions and similar shape, crystals belonged to two different space groups (P1 and P21; Table S1). An initial model was determined by ARCIMBOLDO_LITE (92) in its COILED_COIL mode (93) using a merged P21 dataset assembled from three individual datasets from different crystals, cut to a final resolution of 2.5 Å. ARCIMBOLDO (94) uses PHASER (95, 96) to place individual α-helices by eLLG (expected log likelihood-gain)-guided molecular replacement (97), then expand partial solutions with SHELXE (98) through density modification and autotracing into a complete model (99). Phases from the initial ARCIMBOLDO model (393 residues) were used to identify selenomethionine sites, which were then supplied to the Phenix Autosol module (100) for phase calculation in PHASER (101), density modification including two-fold NCS averaging in RESOLVE (102), and initial model building in RESOLVE (102). Initial models from ARCIMBOLDO and RESOLVE were manually rebuilt in COOT and refined in phenix.refine (103) against a single 2.5 Å-resolution dataset collected from crystals of selenomethionine-substituted protein. The register of all four protein chains in the final model, and their identity as SYCP3^CC^, were verified by anomalous difference maps showing the location of selenomethionine residues. While ARCIMBOLDO successfully determined the structure in the P1 crystal form, the initial P1 model used for rebuilding and refinement was generated by molecular replacement in PHASER using the P21 model. The P1 model was refined against a 2.2 Å-resolution dataset generated by merging five independent datasets collected at APS beamline 24ID-E and SSRL beamline 14-1.

#### Support Statement - Advanced Photon Source NE-CAT beamline 24ID-C

This work is based upon research conducted at the Northeastern Collaborative Access Team beamlines, which are funded by the National Institute of General Medical Sciences from the National Institutes of Health (P41 GM103403). The Pilatus 6M detector on 24-ID-C beam line is funded by a NIH-ORIP HEI grant (S10 RR029205). This research used resources of the Advanced Photon Source, a U.S. Department of Energy (DOE) Office of Science User Facility operated for the DOE Office of Science by Argonne National Laboratory under Contract No. DE-AC02-06CH11357.

#### Support Statement - Stanford Synchrotron Radiation Lightsource beamline 14-1

Use of the Stanford Synchrotron Radiation Lightsource, SLAC National Accelerator Laboratory, is supported by the U.S. Department of Energy, Office of Science, Office of Basic Energy Sciences under Contract No. DE-AC02-76SF00515. The SSRL Structural Molecular Biology Program is supported by the DOE Office of Biological and Environmental Research, and by the National Institutes of Health, National Institute of General Medical Sciences (including P41GM103393). The contents of this publication are solely the responsibility of the authors and do not necessarily represent the official views of NIGMS or NIH.

### Small-angle X-ray Scattering (SAXS)

For SAXS, *Mm* SYCP2^CC^:SYCP3^CC^ was diluted to 1, 3, or 6 mg/mL in a buffer containing 20 mM Tris-HCl pH 8.5, 300 mM NaCl, 2% glycerol, and 1 mM DTT. The *Hs* SYCP3^CC^-SYCP2^CC^ fusion was diluted to 2, 4 or 8 mg/mL in a buffer containing 20 mM Tris-HCl pH 7.5, 300 mM NaCl, and 1 mM DTT. SAXS data were collected at the SIBYLS Beamline 12.3.1 at the Advanced Light Source, Lawrence Berkeley National Lab (which is funded by DOE BER Integrated Diffraction Analysis Technologies (IDAT) program and NIGMS grant P30 GM124169-01, ALS-ENABLE) (104). For each sample, thirty 0.3-second exposures were taken and integrated, for a total exposure time of 10 seconds. Exposures were radially averaged and buffer-subtracted to yield SAXS scattering curves. SAXS data analysis was performed with ScÅtter (https://bl1231.als.lbl.gov/scatter/) and the ATSAS SAXS analysis suite (https://www.embl-hamburg.de/biosaxs/software.html) (104).

### Crosslinking mass spectrometry (XLMS)

For cross-linking of *Hs* SYCP3^CC^-SYCP2^CC^, the protein was diluted to 1 mg/mL in a buffer containing 20 mM HEPES pH 7.5, 300 mM NaCl, 10% glycerol, and 1 mM DTT. Crosslinking was performed by addition of 0.2, 0.5, or 1 mM isotopically-coded D0/D12 BS^3^ (bis-sulfosuccinimidylsuberate; Creative Molecules) for 60 minutes at room temperature. The reaction was quenched by the addition of 100 mM NH^4^HCO^3^ and further incubation at 30°C for 10 minutes. Quenched reactions were supplemented with 8M urea to a final concentration of 6M. Subsequent to reduction and alkylation, crosslinked proteins were digested with Lys-C (1:50 w/w, Wako) for 3 h, diluted with 50 mM ammonium bicarbonate to 1M urea and digested with trypsin (1:50 w/w, Promega) overnight. Crosslinked peptides were purified by reversed phase chromatography using C18 cartridges (Sep-Pak, Waters). Crosslink fractions by peptide size exclusion chromatography and analyzed by tandem mass spectrometry (Orbitrap Elite, Thermo Scientific) (105). Fragment ion spectra were searched and crosslinks identified by the dedicated software program xQuest (106). All unique detected crosslinks are listed in Tables S2 and S3.

### Yeast genetics and imaging

All yeast strains were derived from the SK1-related diploid strain NH144 (Table S4) (107, 108). For *Sc-Zr* Red1 chimeras, a homologous recombination template was generated to replace residues 734-827 with residues 705-798 or 705-791 of *Zr* Red1, followed by a *KanMX* selection marker, and integrated into the *RED1* locus. For expression of Red1^1-362^, a homologous recombination template was generated to replace residues 363-827 with a 6xHis-3xHA tag followed by a *KanMX* selection marker, and integrated into the *RED1* locus. For spore viability, cells were grown on YPD agar, patched onto SPO medium (1% KOAc) for 48-72 hours, then tetrads were dissected onto YPD agar and grown 3 days for analysis.

For synchronous meiosis and fluorescence imaging, cells were grown in YPD, then diluted into BYTA (YEP + 1% KOAc/50 mM potassium phthalate) at OD600 = 0.3, grown overnight, then washed and resuspended in SPO medium (0.3% KOAc [pH 7.0]) at OD600 = 2.0 at 30° C to induce sporulation. Samples were removed hourly for 10 hours, fixed, and stained with DAPI and either rabbit anti-Red1 (a kind gift from Nancy Hollingsworth) or rat anti-HA primary antibodies (Sigma 11867431001) followed by secondary antibodies (Cy3-linked anti-rabbit, Jackson IR 711166152; AlexaFluor 647-linked anti-rat, Jackson IR 712605153). Fluorescence microscopy was performed on a deconvolution microscope (DeltaVision; Applied Precision/GE Healthcare) equipped with a charge-coupled device camera (CoolSNAP; Roper Scientific) controlled by a softWoRx workstation (DeltaVision; Applied Precision/GE Healthcare).

## Acknowledgements

The authors thank the staffs of the Stanford Synchrotron Light Source and the Advanced Photon Source sector 24 for assistance with collecting x-ray diffraction data, the staff of the Advanced Light Source Sibyls Beamline 12.3.1 for assistance with SAXS data collection and processing, and T. Booth at the UCSD Cryo-Electron Microscopy Facility for assistance with electron microscopy. We thank members of the Corbett lab, A. Desai, and Y. Kim for critical reading and helpful discussions. S.U. is supported by the National Science Foundation (Graduate Research Fellowship). I.U. and I.C. are supported by grants BIO2015-64216-P and MDM2014-0435 (the Spanish Ministry of Science, Innovation and Universities). K.D.C. is supported by the Ludwig Institute for Cancer Research and the National Institutes of Health (R01 GM104141). K.D.C. and F.H. acknowledge joint support from the Human Frontiers Science Program (#RGP0008/2015).

## Competing Interests

The authors declare no competing interests.

## References

1. Zickler,D. and Kleckner,N. (1999) Meiotic Chromosomes: Integrating Structure and Function. Annu. Rev. Genet., 33, 603–754.

2. Moens,P.B., Pearlman,R.E., Heng,H.H. and Traut,W. (1998) Chromosome cores and chromatin at meiotic prophase. Curr. Top. Dev. Biol., 37, 241–262.

3. van Heemst,D. and Heyting,C. (2000) Sister chromatid cohesion and recombination in meiosis. Chromosoma, 109, 10–26.

4. Panizza,S., Mendoza,M.A., Berlinger,M., Huang,L., Nicolas,A., Shirahige,K. and Klein,F. (2011) Spo11-accessory proteins link double-strand break sites to the chromosome axis in early meiotic recombination. Cell, 146, 372–383.

5. Zickler,D. and Kleckner,N. (2015) Recombination, Pairing, and Synapsis of Homologs during Meiosis. Cold Spring Harb Perspect Biol, 7, a016626.

6. Kim,K.P., Weiner,B.M., Zhang,L., Jordan,A., Dekker,J. and Kleckner,N. (2010) Sister cohesion and structural axis components mediate homolog bias of meiotic recombination. Cell, 143, 924–937.

7. Subramanian,V.V. and Hochwagen,A. (2014) The meiotic checkpoint network: step-by-step through meiotic prophase. Cold Spring Harb Perspect Biol, 6, a016675–a016675.

8. Lao,J.P. and Hunter,N. (2010) Trying to avoid your sister. PLoSBiol., 8, e1000519.

9. Hollingsworth,N.M. (2010) Phosphorylation and the creation of interhomolog bias during meiosis in yeast. Cell Cycle, 9, 436–437.

10. Humphryes,N. and Hochwagen,A. (2014) A non-sister act: recombination template choice during meiosis. Experimental Cell Research, 329, 53–60.

11. Carballo,J.A., Johnson,A.L., Sedgwick,S.G. and Cha,R.S. (2008) Phosphorylation of the axial element protein Hop1 by Mec1/Tel1 ensures meiotic interhomolog recombination. Cell, 132, 758–770.

12. Niu,H., Li,X., Job,E., Park,C., Moazed,D., Gygi,S.P. and Hollingsworth,N.M. (2007) Mek1 kinase is regulated to suppress double-strand break repair between sister chromatids during budding yeast meiosis. Molecular and cellular biology, 27, 5456–5467.

13. Niu,H., Wan,L., Busygina,V., Kwon,Y., Allen,J.A., Li,X., Kunz,R.C., Kubota,K., Wang,B., Sung,P., et al. (2009) Regulation of meiotic recombination via Mek1-mediated Rad54 phosphorylation. Molecular Cell, 36, 393–404.

14. Lao,J.P., Cloud,V., Huang,C.-C., Grubb,J., Thacker,D., Lee,C.-Y., Dresser,M.E., Hunter,N. and Bishop,D.K. (2013) Meiotic crossover control by concerted action of Rad51-Dmc1 in homolog template bias and robust homeostatic regulation. PLoS Genet, 9, e1003978.

15. Onn,I., Heidinger-Pauli,J.M., Guacci,V., Ünal,E. and Koshland,D.E. (2008) Sister Chromatid Cohesion: A Simple Concept with a Complex Reality. Annu. Rev. Cell Dev. Biol., 24, 105–129.

16. Vader,G. and Musacchio,A. (2014) HORMA Domains at the Heart of Meiotic Chromosome Dynamics. Developmental Cell, 31, 389–391.

17. Aravind,L. and Koonin,E.V. (1998) The HORMA domain: a common structural denominator in mitotic checkpoints, chromosome synapsis and DNA repair. Trends Biochem. Sci., 23, 284–286.

18. Hollingsworth,N.M. and Byers,B. (1989) HOP1: a yeast meiotic pairing gene. Genetics, 121, 445–462.

19. Wojtasz,L., Daniel,K., Roig,I., Bolcun-Filas,E., Xu,H., Boonsanay,V., Eckmann,C.R., Cooke,H.J., Jasin,M., Keeney,S., et al. (2009) Mouse HORMAD1 and HORMAD2, two conserved meiotic chromosomal proteins, are depleted from synapsed chromosome axes with the help of TRIP13 AAA-ATPase. PLoS Genet, 5, e1000702.

20. Zetka,M.C., Kawasaki,I., Strome,S. and Müller,F. (1999) Synapsis and chiasma formation in Caenorhabditis elegans require HIM-3, a meiotic chromosome core component that functions in chromosome segregation. Genes & Development, 13, 2258–2270.

21. Couteau,F. and Zetka,M. (2005) HTP-1 coordinates synaptonemal complex assembly with homolog alignment during meiosis in C. elegans. Genes & Development, 19, 2744–2756.

22. Martinez-Perez,E. and Villeneuve,A.M. (2005) HTP-1-dependent constraints coordinate homolog pairing and synapsis and promote chiasma formation during C. elegans meiosis. Genes & Development, 19, 2727–2743.

23. Lorenz,A., Wells,J.L., Pryce,D.W., Novatchkova,M., Eisenhaber,F., McFarlane,R.J. and Loidl,J. (2004) S. pombe meiotic linear elements contain proteins related to synaptonemal complex components. J Cell Sci, 117, 3343–3351.

24. Caryl,A.P., Armstrong,S.J., Jones,G.H. and Franklin,F.C.H. (2000) A homologue of the yeast HOP1 gene is inactivated in the Arabidopsis meiotic mutant asy1. Chromosoma, 109, 62–71.

25. West,A.M.V., Komives,E.A. and Corbett,K.D. (2018) Conformational dynamics of the Hop1 HORMA domain reveal a common mechanism with the spindle checkpoint protein Mad2. Nucleic Acids Research, 46, 279–292.

26. Woltering,D., Baumgartner,B., Bagchi,S., Larkin,B., Loidl,J., de los Santos,T. and Hollingsworth,N.M. (2000) Meiotic segregation, synapsis, and recombination checkpoint functions require physical interaction between the chromosomal proteins Red1p and Hop1p. Molecular and cellular biology, 20, 6646–6658.

27. Hollingsworth,N.M. and Ponte,L. (1997) Genetic interactions between HOP1, RED1 and MEK1 suggest that MEK1 regulates assembly of axial element components during meiosis in the yeast Saccharomyces cerevisiae. Genetics, 147, 33–42.

28. Yuan,L., Liu,J.-G., Zhao,J., Brundell,E., Daneholt,B. and Höög,C. (2000) The Murine SCP3 Gene Is Required for Synaptonemal Complex Assembly, Chromosome Synapsis, and Male Fertility. Molecular Cell, 5, 73–83.

29. Yuan,L., Liu,J.-G., Hoja,M.-R., Wilbertz,J., Nordqvist,K. and Höög,C. (2002) Female Germ Cell Aneuploidy and Embryo Death in Mice Lacking the Meiosis-Specific Protein SCP3. Science, 296, 1115–1118.

30. Yang,F., Fuente,R.D.L., Leu,N.A., Baumann,C., McLaughlin,K.J. and Wang,P.J. (2006) Mouse SYCP2 is required for synaptonemal complex assembly and chromosomal synapsis during male meiosis. J Cell Biol, 173, 497–507.

31. Pelttari,J., Hoja,M.R., Yuan,L., Liu,J.G., Brundell,E., Moens,P., Santucci-Darmanin,S., Jessberger,R., Barbero,J.L., Heyting,C., et al. (2001) A meiotic chromosomal core consisting of cohesin complex proteins recruits DNA recombination proteins and promotes synapsis in the absence of an axial element in mammalian meiotic cells. Molecular and cellular biology, 21, 5667–5677.

32. Shin,Y.-H., Choi,Y., Erdin,S.U., Yatsenko,S.A., Kloc,M., Yang,F., Wang,P.J., Meistrich,M.L. and Rajkovic,A. (2010) Hormad1 mutation disrupts synaptonemal complex formation, recombination, and chromosome segregation in mammalian meiosis. PLoS Genet, 6, e1001190.

33. Ferdous,M., Higgins,J.D., Osman,K., Lambing,C., Roitinger,E., Mechtler,K., Armstrong,S.J., Perry,R., Pradillo,M., Cuñado,N., et al. (2012) Inter-Homolog Crossing-Over and Synapsis in Arabidopsis Meiosis Are Dependent on the Chromosome Axis Protein AtASY3. PLoS Genet, 8, e1002507.

34. Osman,K., Yang,J., Roitinger,E., Lambing,C., Heckmann,S., Howell,E., Cuacos,M., Imre,R., Dürnberger,G., Mechtler,K., et al. (2017) Affinity proteomics reveals extensive phosphorylation of the Brassica chromosome axis protein ASY1 and a network of associated proteins at prophase I of meiosis. Plant J., 93, 17–33.

35. Offenberg,H.H., Schalk,J.A., Meuwissen,R.L., van Aalderen,M., Kester,H.A., Dietrich,A.J. and Heyting,C. (1998) SCP2: a major protein component of the axial elements of synaptonemal complexes of the rat. Nucleic Acids Research, 26, 2572–2579.

36. Smith,A.V. and Roeder,G.S. (1997) The Yeast Red1 Protein Localizes to the Cores of Meiotic Chromosomes. J Cell Biol, 136, 957–967.

37. Börner,G.V., Barot,A. and Kleckner,N. (2008) Yeast Pch2 promotes domainal axis organization, timely recombination progression, and arrest of defective recombinosomes during meiosis. Proceedings of the National Academy of Sciences, 105, 3327–3332.

38. Joshi,N., Barot,A., Jamison,C. and Börner,G.V. (2009) Pch2 links chromosome axis remodeling at future crossover sites and crossover distribution during yeast meiosis. PLoS Genet, 5, e1000557.

39. Roig,I., Dowdle,J.A., Toth,A., de Rooij,D.G., Jasin,M. and Keeney,S. (2010) Mouse TRIP13/PCH2 Is Required for Recombination and Normal Higher-Order Chromosome Structure during Meiosis. PLoS Genet, 6, e1001062–19.

40. Lambing,C., Osman,K., Nuntasoontorn,K., West,A., Higgins,J.D., Copenhaver,G.P., Yang,J., Armstrong,S.J., Mechtler,K., Roitinger,E., et al. (2015) Arabidopsis PCH2 Mediates Meiotic Chromosome Remodeling and Maturation of Crossovers. PLoS Genet, 11, e1005372.

41. San-Segundo,P.A. and Roeder,G.S. (1999) Pch2 links chromatin silencing to meiotic checkpoint control. 97, 313–324.

42. Vader,G. (2015) Pch2(TRIP13): controlling cell division through regulation of HORMA domains. Chromosoma, 124, 333–339.

43. Page,S.L. and Hawley,R.S. (2004) The genetics and molecular biology of the synaptonemal complex. Annu. Rev. Cell Dev. Biol., 20, 525–558.

44. Dunce,J.M., Dunne,O.M., Ratcliff,M., Millán,C., Madgwick,S., Usón,I. and Davies,O.R. (2018) Structural basis of meiotic chromosome synapsis through SYCP1 self-assembly. Nature Publishing Group, 25, 557–569.

45. Lu,J., Gu,Y., Feng,J., Zhou,W., Yang,X. and Shen,Y. (2014) Structural Insight into the Central Element Assembly of the Synaptonemal Complex. Sci Rep, 4, 128.

46. Cahoon,C.K., Yu,Z., Wang,Y., Guo,F., Unruh,J.R., Slaughter,B.D. and Hawley,R.S. (2017) Superresolution expansion microscopy reveals the three-dimensional organization of the Drosophilasynaptonemal complex. Proceedings of the National Academy of Sciences, 114, E6857–E6866.

47. Köhler,S., Wojcik,M., Xu,K. and Dernburg,A.F. (2017) Superresolution microscopy reveals the three-dimensional organization of meiotic chromosome axes in intact Caenorhabditis eleganstissue. Proceedings of the National Academy of Sciences, 114, E4734–E4743.

48. Schücker,K., Holm,T., Franke,C., Sauer,M. and Benavente,R. (2015) Elucidation of synaptonemal complex organization by super-resolution imaging with isotropic resolution. Proceedings of the National Academy of Sciences, 112, 2029–2033.

49. Syrjänen,J.L., Pellegrini,L. and Davies,O.R. (2014) A molecular model for the role of SYCP3 in meiotic chromosome organisation. eLife, 3, 213–18.

50. Yuan,L., Pelttari,J., Brundell,E., Björkroth,B., Zhao,J., Liu,J.-G., Brismar,H., Daneholt,B. and Höög,C. (1998) The Synaptonemal Complex Protein SCP3 Can Form Multistranded, Cross-striated Fibers In Vivo. J Cell Biol, 142, 331–339.

51. Syrjänen,J.L., Heller,I., Candelli,A., Davies,O.R., Peterman,E.J.G., Wuite,G.J.L. and Pellegrini,L. (2017) Single-molecule observation of DNA compaction by meiotic protein SYCP3. eLife, 6, 595.

52. Bollschweiler,D., Radu,L., Plitzko,J.M., Henderson,R.M., Mela,I. and Pellegrini,L. (2018) Reconstitution of a flexible SYCP3-DNA fibre suggests a mechanism for SYCP3 coating of the meiotic chromosome axis. bioRxiv, 10.1101/369439.

53. Klein,F., Mahr,P., Galova,M., Buonomo,S.B., Michaelis,C., Nairz,K. and Nasmyth,K. (1999) A central role for cohesins in sister chromatid cohesion, formation of axial elements, and recombination during yeast meiosis. 98, 91–103.

54. Eichinger,C.S. and Jentsch,S. (2010) Synaptonemal complex formation and meiotic checkpoint signaling are linked to the lateral element protein Red1. Proceedings of the National Academy of Sciences, 107, 11370–11375.

55. Lin,F.-M., Lai,Y.-J., Shen,H.-J., Cheng,Y.-H. and Wang,T.-F. (2010) Yeast axial-element protein, Red1, binds SUMO chains to promote meiotic interhomologue recombination and chromosome synapsis. EMBO J., 29, 586–596.

56. Rockmill,B. and Roeder,G.S. (1988) RED1: a yeast gene required for the segregation of chromosomes during the reductional division of meiosis. Proceedings of the National Academy of Sciences, 85, 6057–6061.

57. Rockmill,B. and Roeder,G.S. (1990) Meiosis in asynaptic yeast. Genetics, 126, 563–574.

58. Biswas,U., Hempel,K., Llano,E., Pendas,A. and Jessberger,R. (2016) Distinct Roles of Meiosis-Specific Cohesin Complexes in Mammalian Spermatogenesis. PLoS Genet, 12, e1006389.

59. Ward,A., Hopkins,J., Mckay,M., Murray,S. and Jordan,P.W. (2016) Genetic Interactions Between the Meiosis-Specific Cohesin Components, STAG3, REC8, and RAD21L. G3: Genes, Genomes, Genetics, 6, 1713–1724.

60. Winters,T., McNicoll,F. and Jessberger,R. (2014) Meiotic cohesin STAG3 is required for chromosome axis formation and sister chromatid cohesion. EMBO J., 33, 1256–1270.

61. Fukuda,T., Fukuda,N., Agostinho,A., Hernández,A.H., Kouznetsova,A. and Höög,C. (2014) STAG3-mediated stabilization of REC8 cohesin complexes promotes chromosome synapsis during meiosis. EMBO J., 33, 1243–1255.

62. Fukuda,T., Daniel,K., Wojtasz,L., Toth,A. and Höög,C. (2010) A novel mammalian HORMA domain-containing protein, HORMAD1, preferentially associates with unsynapsed meiotic chromosomes. Experimental Cell Research, 316, 158–171.

63. Kogo,H., Tsutsumi,M., Inagaki,H., Ohye,T., Kiyonari,H. and Kurahashi,H. (2012) HORMAD2 is essential for synapsis surveillance during meiotic prophase via the recruitment of ATR activity. Genes Cells, 17, 897–912.

64. Li,X.C., Bolcun-Filas,E. and Schimenti,J.C. (2011) Genetic Evidence That Synaptonemal Complex Axial Elements Govern Recombination Pathway Choice in Mice. Genetics, 189, 71–82.

65. Winkel,K., Alsheimer,M., Öllinger,R. and Benavente,R. (2008) Protein SYCP2 provides a link between transverse filaments and lateral elements of mammalian synaptonemal complexes. Chromosoma, 118, 259–267.

66. Llano,E., Herrán,Y., García-Tuñón,I., Gutiérrez-Caballero,C., de Álava,E., Barbero,J.L., Schimenti,J., de Rooij,D.G., Sánchez-Martín,M. and Pendás,A.M. (2012) Meiotic cohesin complexes are essential for the formation of the axial element in mice. The Journal of Cell Biology, 197, 877–885.

67. Kim,Y., Rosenberg,S.C., Kugel,C.L., Kostow,N., Rog,O., Davydov,V., Su,T.Y., Dernburg,A.F. and Corbett,K.D. (2014) The chromosome axis controls meiotic events through a hierarchical assembly of HORMA domain proteins. Developmental Cell, 31, 487–502.

68. Feng,J., Fu,S., Cao,X., Wu,H., Lu,J., Zeng,M., Liu,L., Yang,X. and Shen,Y. (2017) Synaptonemal complex protein 2 (SYCP2) mediates the association of the centromere with the synaptonemal complex. Protein Cell, 8, 538–543.

69. Tarsounas,M., Pearlman,R.E., Gasser,P.J., Park,M.S. and Moens,P.B. (1997) Protein-protein interactions in the synaptonemal complex. Mol. Biol. Cell, 8, 1405–1414.

70. Yang,F., La Fuente,De,R., Leu,N.A., Baumann,C., McLaughlin,K.J. and Wang,P.J. (2006) Mouse SYCP2 is required for synaptonemal complex assembly and chromosomal synapsis during male meiosis. J Cell Biol, 173, 497–507.

71. Baier,A., Alsheimer,M., Volff,J.N. and Benavente,R. (2007) Synaptonemal Complex Protein SYCP3 of the Rat: Evolutionarily Conserved Domains and the Assembly of Higher Order Structures. Sex Dev, 1, 161–168.

72. Feigin,L.A. and Svergun,D.I. (1987) Structure Analysis by Small-Angle X-Ray and Neutron Scattering. Springer US.

73. Glatter,O. and Kratky,O. (1982) Small Angle X-ray Scattering. Academic Press.

74. Zamariola,L., Tiang,C.L., De Storme,N., Pawlowski,W. and Geelen,D. (2014) Chromosome segregation in plant meiosis. Front. Plant Sci., 5, 135.

75. Lam,W.S. (2005) Characterization of Arabidopsis thaliana SMC1 and SMC3: evidence that AtSMC3 may function beyond chromosome cohesion. J Cell Sci, 118, 3037–3048.

76. Cai,X., Dong,F., Edelmann,R.E. and Makaroff,C.A. (2003) The Arabidopsis SYN1 cohesin protein is required for sister chromatid arm cohesion and homologous chromosome pairing. J Cell Sci, 116, 2999–3007.

77. Bhatt,A.M., Lister,C., Page,T., Fransz,P., Findlay,K., Jones,G.H., Dickinson,H.G. and Dean,C. (1999) The DIF1 gene of Arabidopsis is required for meiotic chromosome segregation and belongs to the REC8/RAD21 cohesin gene family. Plant J., 19, 463–472.

78. Armstrong,S.J., Caryl,A.P., Jones,G.H. and Franklin,F.C.H. (2002) Asy1, a protein required for meiotic chromosome synapsis, localizes to axis-associated chromatin in Arabidopsis and Brassica. J Cell Sci, 115, 3645–3655.

79. Chambon,A., West,A., Vezon,D., Horlow,C., De Muyt,A., Chelysheva,L., Ronceret,A., Darbyshire,A.R., Osman,K., Heckmann,S., et al. (2018) Identification of ASYNAPTIC4, a component of the meiotic chromosome axis. Plant Physiology, 10.1104/pp.17.01725.

80. Novak,I., Wang,H., Revenkova,E., Jessberger,R., Scherthan,H. and Höög,C. (2008) Cohesin Smc1β determines meiotic chromatin axis loop organization. J Cell Biol, 180, 83–90.

81. Revenkova,E., Eijpe,M., Heyting,C., Hodges,C.A., Hunt,P.A., Liebe,B., Scherthan,H. and Jessberger,R. (2004) Cohesin SMC1β is required for meiotic chromosome dynamics, sister chromatid cohesion and DNA recombination. Nat. Cell Biol., 6, 555–562.

82. Bannister,L.A., Reinholdt,L.G., Munroe,R.J. and Schimenti,J.C. (2004) Positional cloning and characterization of mouse mei8, a disrupted allelle of the meiotic cohesin Rec8. Genesis, 40, 184–194.

83. Xu,H., Beasley,M.D., Warren,W.D., van der Horst,G.T.J. and McKay,M.J. (2005) Absence of mouse REC8 cohesin promotes synapsis of sister chromatids in meiosis. Developmental Cell, 8, 949–961.

84. Herrán,Y., Caballero,C.G., Martín,M.S., Hernández,T., Viera,A., Barbero,J.L., de Álava,E., de Rooij,D.G., Suja,J.Á., Llano,E., et al. (2011) The cohesin subunit RAD21L functions in meiotic synapsis and exhibits sexual dimorphism in fertility. EMBO J., 30, 3091–3105.

85. Hopkins,J., Hwang,G., Jacob,J., Sapp,N., Bedigian,R., Oka,K., Overbeek,P., Murray,S. and Jordan,P.W. (2014) Meiosis-Specific Cohesin Component, Stag3 Is Essential for Maintaining Centromere Chromatid Cohesion, and Required for DNA Repair and Synapsis between Homologous Chromosomes. PLoS Genet, 10, e1004413.

86. Caburet,S., Arboleda,V.A., Llano,E., Overbeek,P.A., Barbero,J.L., Oka,K., Harrison,W., Vaiman,D., Ben-Neriah,Z., García-Tuñón,I., et al. (2014) Mutant Cohesin in Premature Ovarian Failure. NEngl J Med, 370, 943–949.

87. Llano,E., Gomez-H,L., García-Tuñón,I., Sánchez-Martín,M., Caburet,S., Barbero,J.L., Schimenti,J.C., Veitia,R.A. and Pendás,A.M. (2014) STAG3 is a strong candidate gene for male infertility. Hum Mol Genet, 23, 3421–3431.

88. Moses,M.J., Dresser,M.E. and Poorman,P.A. (1984) Composition and role of the synaptonemal complex. Symp. Soc. Exp. Biol., 38, 245–270.

89. Ortiz,R., Echeverría,O.M., Ubaldo,E., Carlos,A., Scassellati,C. and Vázquez-Nin,G.H. (2002) Cytochemical study of the distribution of RNA and DNA in the synaptonemal complex of guinea-pig and rat spermatocytes. Eur JHistochem, 46, 133–142.

90. Rong,M., Matsuda,A., Hiraoka,Y. and Lee,J. (2016) Meiotic cohesin subunits RAD21L and REC8 are positioned at distinct regions between lateral elements and transverse filaments in the synaptonemal complex of mouse spermatocytes. J. Reprod. Dev., 62, 623–630.

91. Sun,X., Huang,L., Markowitz,T.E., Blitzblau,H.G., Chen,D., Klein,F. and Hochwagen,A. (2015) Transcription dynamically patterns the meiotic chromosome-axis interface. eLife, 4, 8522.

92. Sammito,M., Millán,C., Frieske,D., Rodríguez-Freire,E., Borges,R.J. and Usón,I. (2015) ARCIMBOLDO_LITE: single-workstation implementation and use. Acta Crystallogr D Biol Crystallogr, 71, 1921–1930.

93. Caballero,I., Sammito,M., Millán,C., Lebedev,A., Soler,N. and Usón,I. (2018) ARCIMBOLDO on coiled coils. Acta Crystallogr D Struct Biol, 74, 194–204.

94. Millán,C., Sammito,M. and Usón,I. (2015) Macromolecular ab initio phasing enforcing secondary and tertiary structure. IUCr J, 2, 95–105.

95. McCoy,A.J., Grosse-Kunstleve,R.W., Adams,P.D., Winn,M.D., Storoni,L.C. and Read,R.J. (2007) Phaser crystallographic software. JAppl Crystallogr, 40, 658–674.

96. Read,R.J. and McCoy,A.J. (2016) A log-likelihood-gain intensity target for crystallographic phasing that accounts for experimental error. Acta Crystallogr D Struct Biol, 72, 375–387.

97. Oeffner,R.D., Afonine,P.V., Millán,C., Sammito,M., Usón,I., Read,R.J. and McCoy,A.J. (2018) On the application of the expected log-likelihood gain to decision making in molecular replacement. - PubMed - NCBI. Acta Crystallogr D Struct Biol, 74, 245–255.

98. Usón,I. and Sheldrick,G.M. (2018) An introduction to experimental phasing of macromolecules illustrated by SHELX; new autotracing features. Acta Crystallogr D Struct Biol, 74, 106–116.

99. Usón,I., Stevenson,C.E.M., Lawson,D.M. and Sheldrick,G.M. (2007) Structure determination of the O-methyltransferase NovP using the ‘free lunch algorithm’ as implemented in SHELXE. Acta Crystallogr D Biol Crystallogr, 63, 1069–1074.

100. Terwilliger,T.C., Adams,P.D., Read,R.J., McCoy,A.J., Moriarty,N.W., Grosse-Kunstleve,R.W., Afonine,P.V., Zwart,P.H., Hung,L.W.IUCr (2009) Decision-making in structure solution using Bayesian estimates of map quality: the PHENIX AutoSol wizard. Acta Crystallogr D Biol Crystallogr, 65, 582–601.

101. Read,R.J. and McCoy,A.J. (2011) Using SAD data in Phaser. Acta Crystallogr D Biol Crystallogr, 67, 338–344.

102. Terwilliger,T.C. (2003) SOLVE and RESOLVE: Automated Structure Solution and Density Modification. Methods in Enzymology, 374, 22–37.

103. Adams,P.D., Afonine,P.V., Bunkóczi,G., Chen,V.B., Davis,I.W., Echols,N., Headd,J.J., Hung,L.-W., Kapral,G.J., Grosse-Kunstleve,R.W., et al. (2010) PHENIX: a comprehensive Python-based system for macromolecular structure solution. Acta Crystallogr D Biol Crystallogr, 66, 213–221.

104. Dyer,K.N., Hammel,M., Rambo,R.P., Tsutakawa,S.E., Rodic,I., Classen,S., Tainer,J.A. and Hura,G.L. (2014) High-Throughput SAXS for the Characterization of Biomolecules in Solution: A Practical Approach. In Structural Genomics, Methods in Molecular Biology. Humana Press, Totowa, NJ, Totowa, NJ, Vol. 1091, pp. 245–258.

105. Herzog,F., Kahraman,A., Boehringer,D., Mak,R., Bracher,A., Walzthoeni,T., Leitner,A., Beck,M., Hartl,F.-U., Ban,N., et al. (2012) Structural Probing of a Protein Phosphatase 2A Network by Chemical Cross-Linking and Mass Spectrometry. Science, 337, 1348–1352.

106. Walzthoeni,T., Joachimiak,L.A., Rosenberger,G., Röst,H.L., Malmström,L., Leitner,A., Frydman,J. and Aebersold,R. (2015) xTract: software for characterizing conformational changes of protein complexes by quantitative cross-linking mass spectrometry. Nature methods, 12, 1185–1190.

107. de los Santos,T. and Hollingsworth,N.M. (1999) Red1p, a MEK1-dependent phosphoprotein that physically interacts with Hop1p during meiosis in yeast. J. Biol. Chem., 274, 1783–1790.

108. Hollingsworth,N.M., Ponte,L. and Halsey,C. (1995) MSH5, a novel MutS homolog, facilitates meiotic reciprocal recombination between homologs in Saccharomyces cerevisiae but not mismatch repair. Genes & Development, 9, 1728–1739.

